# ASD mutation of Katnal2 impairs ependymal ciliary motion and causes hydrocephalus

**DOI:** 10.1101/2023.07.03.547302

**Authors:** Shuai Qiu, Huaqiang Cao, Yao Xue, Wenlong Zhao, Shuangyi Xie, Nan Wu, Meilin Song, Yi-Hsuan Pan, Peter W. Baas, Ji Ma, Xiao-Bing Yuan

**Author notes:** These authors jointly supervised this work: Xiao-Bing Yuan, Ji Ma.

## Abstract

Katanin catalytic subunit A1 like 2 (*KATNAL2*) is a high-risk gene associated with autism spectrum disorders (ASD), however its impact on brain development and disease remains unclear. The present study revealed an unexpected role of KATNAL2 in regulating ependymal ciliary motion and cerebrospinal fluid flow during brain development, an important contributing factor for ASD. We discovered a distinct expression pattern of *KATNAL2* in multiciliated ependymal cells of both human and mouse brains. Notably, an ASD-associated mutation of *Katnal2* disrupted its molecular function and resulted in ASD-related behavioral deficits in mice. Additionally, this mutation affected the polarized organization and beating of ependymal cilia, leading to delayed cerebrospinal fluid flow and sustained ventricular enlargement from the early postnatal stage. Conditional ablation of *Katnal2* specifically in the ependymal cells of neonatal mice is sufficient to cause ventricular dilation, whereas no such effect was observed in adult mice. Our findings highlight the importance of ependymal motile cilia and hydrocephalus in ASD, offering insights into its pathogenesis and potential intervention.

## Main

Autism spectrum disorder (ASD) is a complex neurodevelopmental condition characterized by social dysfunction, language impairments, stereotyped or repetitive behaviors, and sensory and motor deficits^1^. Both genetic and environmental factors contribute to the etiology of ASD^1–3^. However, the precise neurodevelopmental mechanisms underlying ASD remain elusive. Remarkably, MRI studies have reported enlarged brain ventricles in individuals with ASD^4–7^, suggesting a potential association between hydrocephalus and ASD.

Ventricular dilation and hydrocephalus can arise from various mechanisms, including obstruction of cerebrospinal fluid (CSF) flow, reduced CSF absorption at arachnoid granulations, and excessive CSF secretion from the choroid plexus (ChP)^8^. Malformation of the ependymal epithelium and dyskinesia of ependymal cilia are also major causes of communicating hydrocephalus^8–10^. Furthermore, defective neurogenesis has been implicated in congenital hydrocephalus^10–12^. Genetic factors play a fundamental role in congenital hydrocephalus, while pathological conditions, such as brain tumors, strokes, infections, and intraventricular hemorrhage, can lead to acquired hydrocephalus^9^. Severe hydrocephalus can damage brain tissues and cause a wide range of physical, behavioral, and cognitive symptoms^8^.

Katanin is a microtubule-severing enzyme composed of a 60 kDa catalytic subunit (katanin-p60) and an 80 kDa B-regulatory subunit (katanin-p80), encoded by the *KATNA1* and *KATNB1* gene, respectively^13, 14^. Katanin family enzymes are essential for cellular processes that depend on microtubule dynamics^15, 16^. Katnal2 is a newly identified member of the Katanin family. The link between *KATNAL2* and ASD was initially discovered through whole exome sequencing of well-diagnosed individuals with ASD to identify *de novo* mutations^17, 18^. Subsequent investigations identified several inherited variants of *KATNAL2* in affected individuals, further supporting its involvement in ASD^19^. Moreover, meta-analysis and genome-wide association studies implicated *KATNAL2* as a novel gene affecting consciousness and divergent thinking^20, 21^. In Xenopus and zebrafish models, Katnal2 was reported to play a crucial role in ciliogenesis, cell mitosis, and brain development^22, 23^. Additionally, Katnal2 is essential for properly forming the microtubule-based axoneme structure during spermatogenesis^24^. Nevertheless, the expression and function of Katnal2 in mammalian brains have yet to be investigated, and it remains unclear how its mutation leads to the autistic phenotype.

## *Katnal2* is expressed by neuro-glial progenitors and ependymal cells

We examined the spatio-temporal correlation between *KATNAL2* and each gene in the entire genome during development based on the BrainSpan human brain transcriptome dataset (www.brainspan.org, RNA-Seq Gencode v10). Gene ontology (GO) analysis of the top 1000 co-expression partners of *KATNAL2* revealed biological processes closely related to the assembly, motility, and functions of motile cilia instead of primary cilia (Fig. 1a and Extended Data Table 1). This suggests that *KATNAL2* is consistently expressed in ependymal cells (EpCs), the only cell type growing motile cilia in the brain, throughout a wide developmental period.

**Fig. 1:**
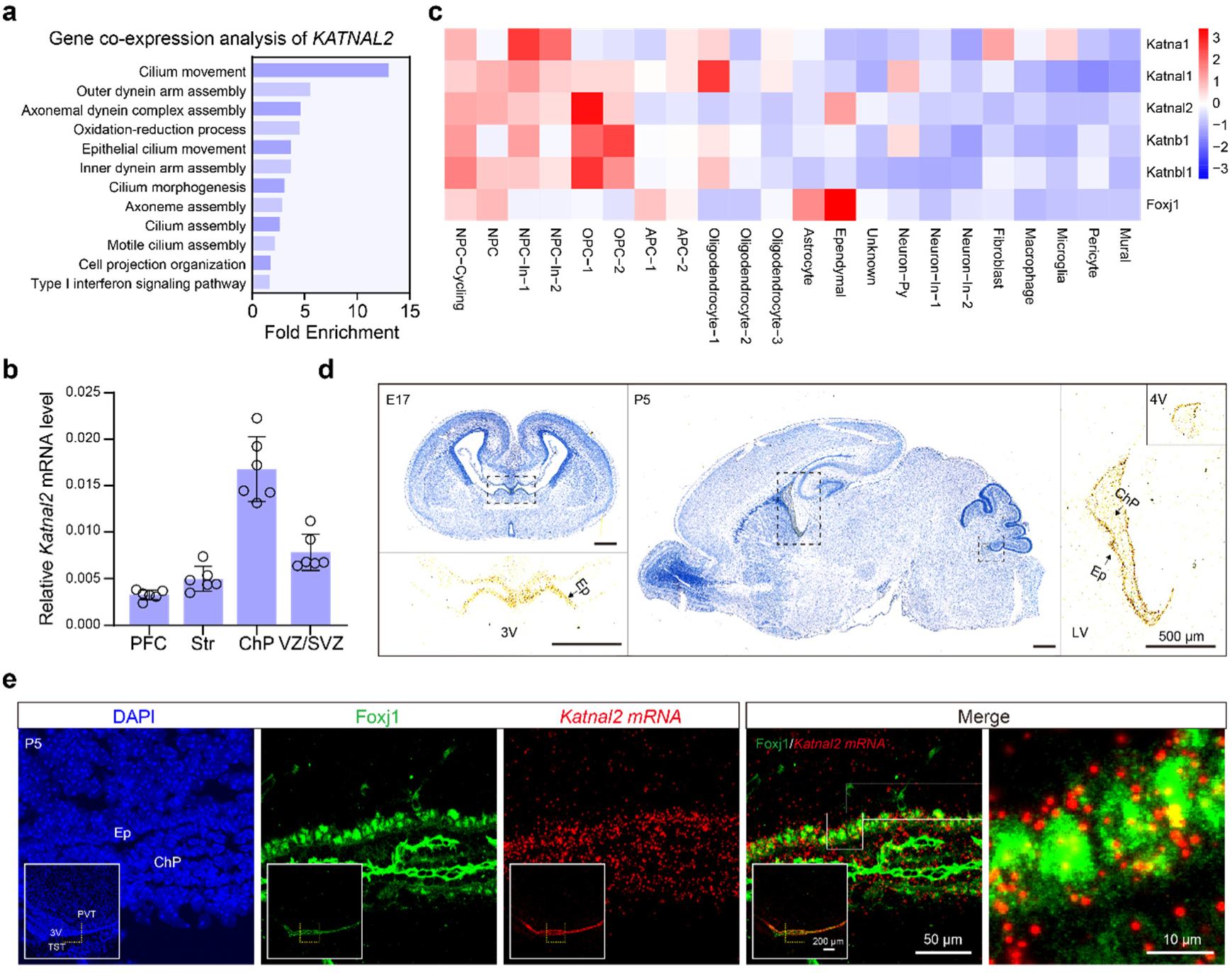
Expression of *Katnal2* in brain ependymal cells. **a**, Gene ontology (GO) analyses of the co-expression partners of *KATNAL2* in developing human brains. GO terms with a Benjamini-Hochberg adjusted *P*-value < 0.05 are shown. **b**, qPCR analysis of *Katnal2* mRNAs at different brain regions of adult mice. **c**, Heat-map presentation of relative expression levels of different *Katanin* family members and *Foxj1* across different cell clusters in postnatal 5 (P5) mouse cortex based on scRNA-seq. The normalized expression of each gene across different cell clusters was the Z-score coded by pseudo-color. **d**, *In-situ* hybridization of *Katnal2* in embryonic day 17 (E17) and P5 mouse brains. Brain sections were Hematoxylin counterstained (blue) to show different brain regions. The high magnification images show *in-situ* hybridization signals of the selected region (dashed line square) of each brain section before Hematoxylin staining. *Katnal2* mRNA (brown signal, arrows) is distributed along the ventricle wall and in the choroid plexus (ChP) of the lateral ventricle (LV), third ventricle (3V), and fourth ventricle (4V). **e**, Expression of *Katnal2* mRNA (red) in Foxj1^+^ (green) cells along the ependymal layer revealed by fluorescent *in-situ* hybridization followed by immunofluorescent staining of Foxj1. The high-magnification images are from the selected region of the low-magnification image of the brain slice shown in the inserts. Ep, ependyma; PVT, paraventricular thalamic nucleus; TST, tectospinal tract.

A previous single-cell RNA sequencing (scRNA-seq) study of developing mouse brains (http://zylkalab.org/datamousecortex)^25^ showed that *katnal2* is enriched in ChP cells at embryonic day 14.5 (E14.5) and at postnatal day 0 (P0), while being expressed at a lower level in radial glial progenitors and immature astrocyte progenitors (Extended Data Fig. 1a). As the ependymal epithelium and the ChP epithelium share the same cell lineage originating from the terminal differentiation of radial glial cells, the results from this scRNA-seq study suggest that *Katnal2* is enriched in the EpC lineage in mouse brains at E14.5 and P0. Our quantitative RT-PCR (qPCR) revealed higher levels of *Katnal2* in the ChP and the ventricular-subventricular zone (VZ/SVZ) than in the gray matter of cortex in 2-month-old mice (Fig. 1b), consistent with a high expression in the EpC/ChP lineage. We further conducted a scRNA-seq analysis of mouse cortex at P5 and found that *Katanin* family members, including *Katna1*, *Katnal1*, *Katnal2*, *Katnb1*, and *Katnbl1*, have a common trend of a higher expression in neural progenitor cells (NPCs) of both the pyramidal lineage and the GABAergic interneuronal lineage and a relatively lower expression in neurons and glial cells (Fig. 1c and Extended Data Table 3). Unlike other *Katanin* family members, *Katnal2* is also expressed in EpCs and in a subgroup of oligodendrocyte progenitor cells (OPCs) at P5 (Fig. 1c). The scRNA-seq also showed that the expression level of *Katnal2* is much lower compared to other *Katanin* family members at P5 (Extended Data Fig. 1b).

We next validated the expression of *Katnal2* in developing and mature mouse brains using *in-situ* hybridization, and found the signal was absent in *Katnal2*-null mice (Extended Fig. 1c). The signal was observed along the ventricle wall, most likely in EpCs and progenitor cells at the VZ/SVZ, as well as in the ChP of all four (lateral, third and fourth) brain ventricles at different developmental stages (Fig. 1d and Extended Data Fig. 1d). We further confirmed the expression of *Katnal2* in EpCs of P5 mouse brains through fluorescent *in-situ* hybridization followed by immunofluorescent staining of Foxj1 (Fig. 1e).

## ASD-associated mutation of *Katnal2* disrupts its molecular function

Multiple disease-related mutations have been identified in the protein-coding region of *KATNAL2* (Extended Data Table 2), including a *de novo* splice site variant (c.510+1G>A) discovered in an autistic child from a Simon Simplex Family^17^. To study the functional consequences of this variant, we used CRISPR/Cas9 gene editing to introduce the same mutation in the mouse *Katnal2* gene (Fig. 2a). As expected, this mutation led to an error in mRNA splicing, resulting in the loss of exon 8 in *Katnal2* transcripts, as confirmed by RT-PCR and Sanger sequencing (Fig. 2b, c). Additionally, during the generation of this splice site mutant allele, we obtained a mutant allele with a large chromosomal segment deletion including exon 8 of *Katnal2* gene (*ΔE8*) (Extended Data Fig. 2a, b). Furthermore, we obtained a *Katnal2*-null (knockout, KO) allele, in which the exon 2-12 region was removed, and a premature stop codon was introduced before exon 13 (Fig. 2a).

**Fig. 2:**
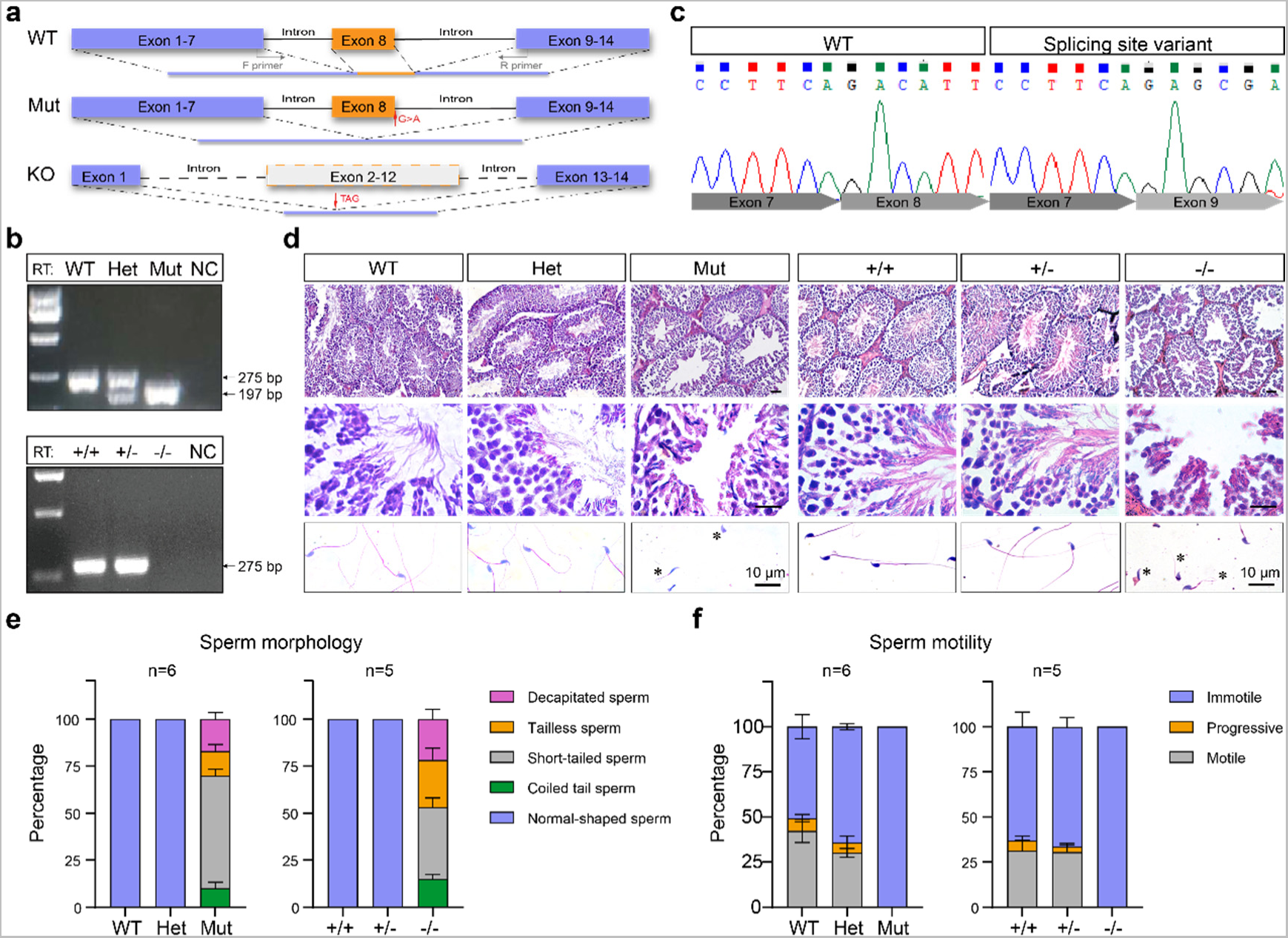
The ASD-associated mutation of Katnal2 disrupts its molecular function. **a**, Schematics of the mutant (Mut) and knockout (KO) alleles of mouse *Katnal2*. Black arrows indicate the location of primers for RT-PCR. **b**, Validation of mutation and knockout of *Katnal2* by RT-PCR of brain mRNAs. The RT-PCR product of the Mut allele was about 78 bp smaller than that of the WT allele (arrows, WT:275 bp; Mut: 195 bp), corresponding to the length of exon 8. There was no RT-PCR product from KO mice. NC, negative control. **c**, Sequencing validation of exon 8 deletion in mRNAs from Mut mice. **d**, Hematoxylin-eosin-stained testis sections (upper and middle) and Giemsa-stained cauda epididymal sperms (lower) from Mut and KO mice compared to that from WT and Het littermates. Asterisks indicate abnormal morphology of cauda epididymal sperms from Mut and KO mice. **e**, Percentages of sperms of different morphological features in mice of different genotypes. **f**, Quantitative analysis of sperm motility of different genotypes. Note the complete loss of sperm motility in Mut and KO mice. Error bars are s.e.m.

For phenotypic analysis, heterozygous *Katnal2* mutant (Het) mice were intercrossed to produce WT, Het, and homozygous mutant (Mut) progeny. Male Mut mice appeared normal in body morphology but were sterile, showing a complete absence of sperm flagella in their testes (Fig. 2d). In contrast, Het mice were largely unaffected. The epididymal sperm content of Mut mice was markedly reduced compared to WT and Het littermates, and the sperm in their cauda epididymis exhibited abnormal head shapes or had truncated or absent tails, resulting in a complete loss of swimming ability (Fig. 2d-f). Similar defects in spermatogenesis and male infertility were observed in homozygous *Katnal2*-null mice (*Katnal2*^-/-^) (Fig. 2d-f), consistent with previous findings^24^. Additionally, male *Katnal2 ΔE8* mice (*E8*^-/-^) were also sterile and displayed similar sperm morphology and motility defects (Extended Data Fig. 2c, d). These results confirm the crucial role of Katnal2 in spermatogenesis and establish that the *de novo* splicing variant of *KATNAL2* found in the autistic child represents a loss-of-function mutation caused by the deletion of exon 8.

## ASD-relevant behavioral deficits in *Katnal2* mutant mice

At 2 to 4 months of age, *Katnal2* mutant mice displayed sex-dependent deficits in several ASD-relevant behavioral paradigms. Both male and female Mut mice exhibited normal social preference in the standard 3-chamber sociability test (stage 2, Fig. 3b, d). However, male Mut mice showed deficits in the social novelty preference (stage 3, Fig. 3b, d). In a resident-intruder experiment, both male and female Mut mice displayed typical defensive responses (data not shown), but female Mut mice exhibited reduced sociability, as indicated by significantly shorter sniffing time towards the intruder (Fig. 3e, f). Female Mut mice also showed deficits in motor learning, as their performance in the rotarod test did not significantly improve after 7 days of training (Fig. 3g, h). Male Mut mice had weaker gripping strength, demonstrated by a shorter duration of holding onto an inverted wire mesh (Fig. 3i, j). In the balance beam test, male Mut mice took significantly longer time to walk through an ascending balance beam (Fig. 3k, l). Both male and female Mut mice walked less smoothly, exhibiting significantly more frequent foot slips and stops than WT littermates (Fig. 3l). These findings align with the high prevalence of motor challenges in children with ASD^26^ and the motor deficits observed in patients with genetic variants in *KATNAL2*^27^. Other behavioral tests, including the open field test, elevated plus-maze test, and marble burying test, did not show significant differences between WT and Mut mice (Fig. 3m-o).

**Fig. 3:**
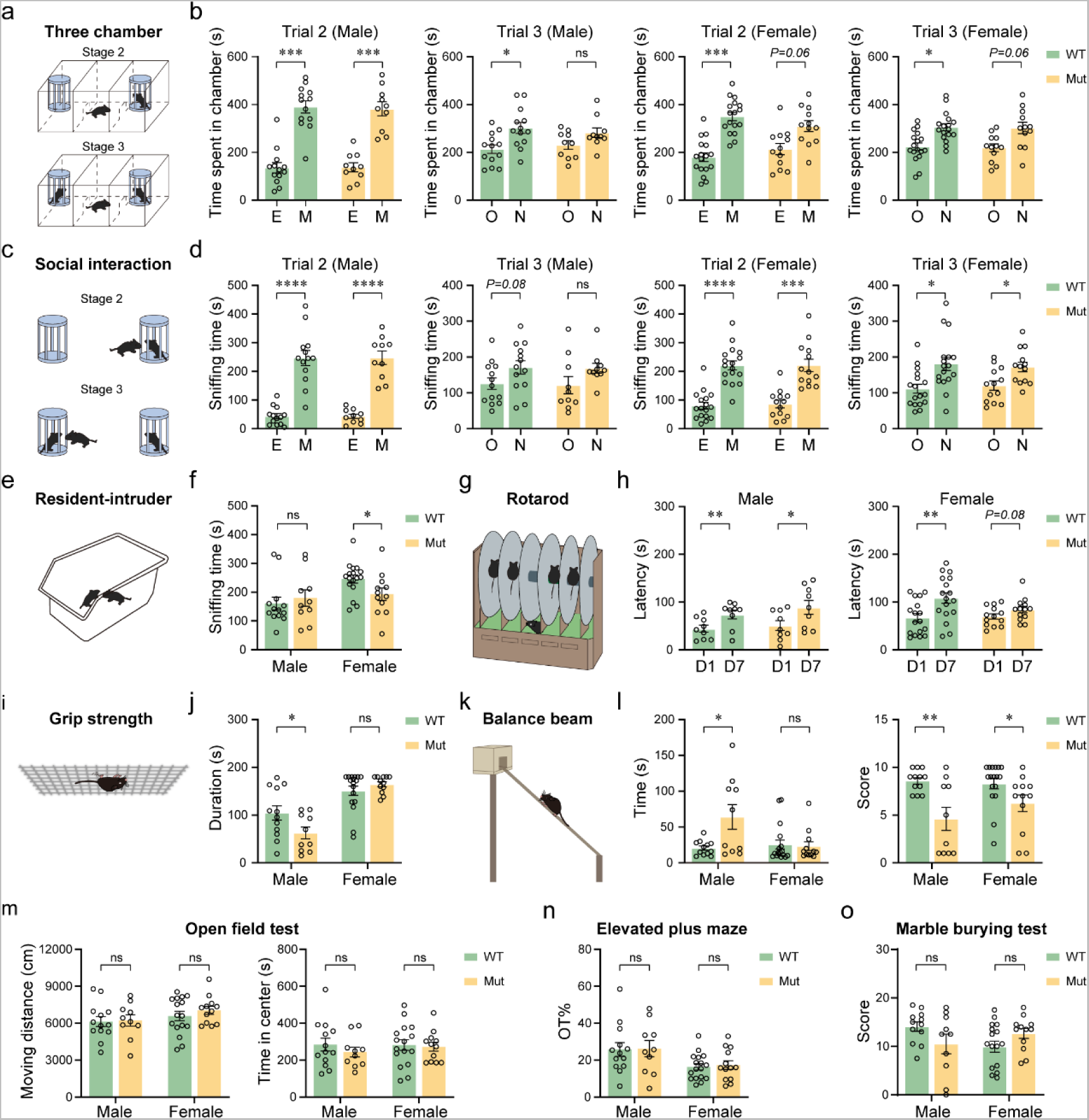
Social and motor deficits in *Katnal2* mutant mice. **a**, Schematics of the standard three-chamber sociability test by measuring the time that the test mouse spent in each side chamber. **b**, Results from social preference and social novelty preference tests by measuring the time spent in each chamber. Male Mut mice showed deficits in the social novelty preference in stage 3. E: Empty, M: mouse, O: familiar mouse, N: novel mouse. **c**, Schematics of the sociability test by measuring the sniffing time of test mouse on stranger mice or object in three-chamber test. **d**, Results from social preference and social novelty preference tests by measuring the sniffing time. **e**, Schematics of the resident-intruder test. **f**, The sniffing time of the resident mouse on the intruder in the resident-intruder test. **g**, Schematics of the rotarod test. **h**, Latency for mice keeping balance on the rotarod. **i**, Schematics of the grip strength test. **j**, Duration for the animal holding to the inverted wire mesh. **k**, Schematics of the balance beam test. **l**, Latency for the animal to reach the goal and the smoothness score based on the count of slippery and stops in the balance beam test. **m**, Total locomotion distance and the time exploring the central zone of mice in the open field test (OFT). **n**, Percentage of time exploring the open arm (OT%) of mice in the elevated plus maze test. **o**, Results of the marble-burying test; n = 10 and 12 for male and female Mut mice, respectively; n = 13 and 17 for male and female WT mice, respectively. Data in histograms are presented as mean ± s.e.m. Each data point (circles) represents the result from one animal. * *P*<0.05, ** *P*<0.01, *** *P*< 0.001, ns: not significant; Student’s *t*-test.

## Brain ventricle enlargement in *Katnal2* mutant mice

At 2 months of age, mice of different genotypes (WT, Het, Mut) had similar body weights and brain sizes (Extended Data Fig. 3a, e-h). However, analysis of Nissl-stained serial coronal sections revealed a significant enlargement in both the lateral ventricle (LV) and dorsal third ventricle (D3V) of Mut mice compared to WT and Het littermates (Fig. 4a-e). The cortex thickness and lamination were not significantly different among genotypes (Extended Data Fig. 3b, c). There was no significant enlargement in the aqueduct (Aq) and the fourth ventricle (4V) of Mut mice (Fig. 4g, h). The enlargement of LVs in Mut mice began during the first postnatal week and persisted until 9 months of age (Fig. 4f). Similar LV enlargement was observed in homozygous *Katnal2*-null mice (Fig. 4c-e) and *ΔE8* mice (Extended Data Fig. 2e), regardless of gender (Fig. 4i, j).

**Fig. 4:**
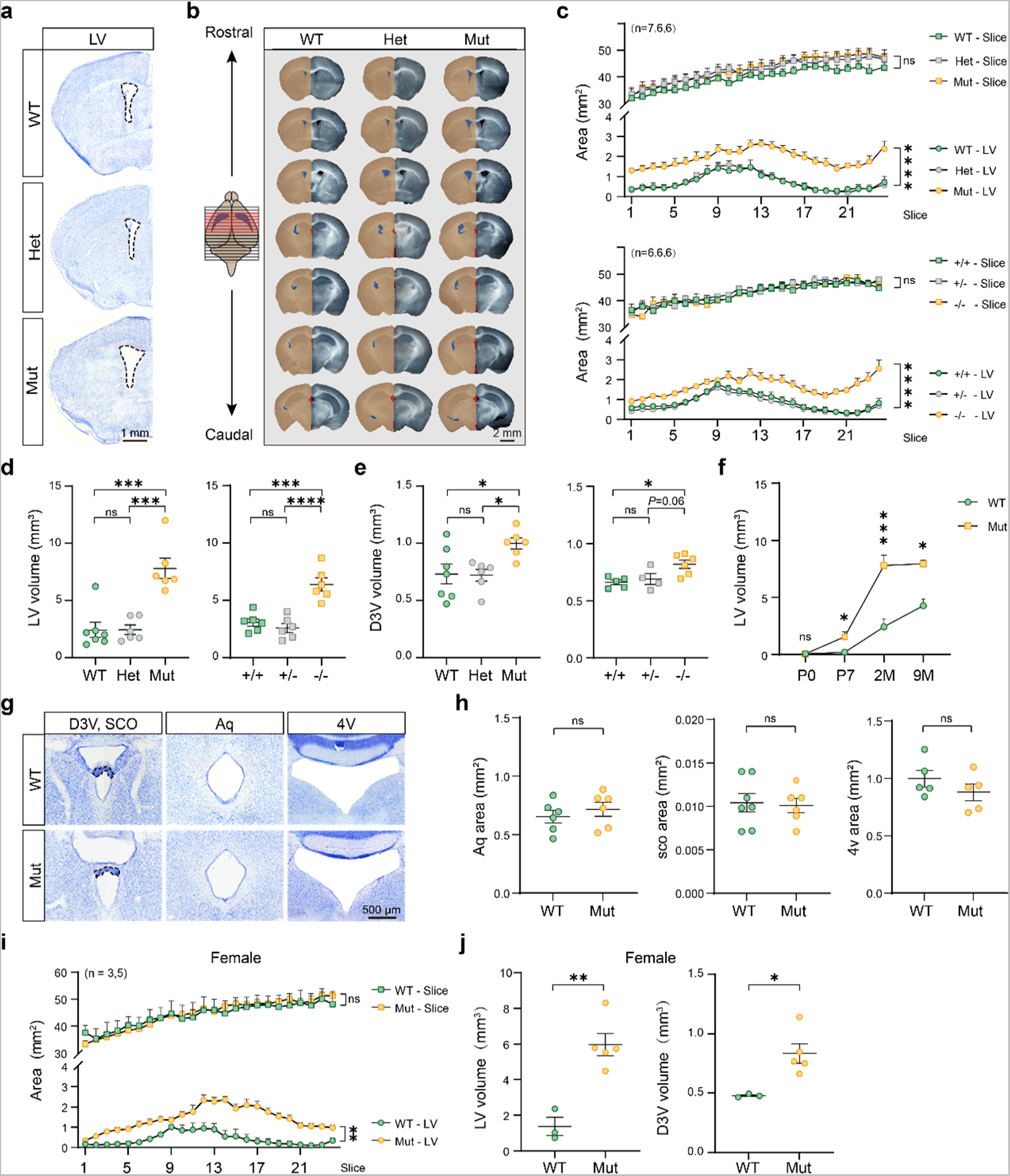
Brain ventricle enlargement in *Katnal2* mutant mice. **a**, Nissl-stained coronal brain sections from 2-month-old (2M) male mice of different genotypes. Note the enlargement of the lateral ventricle (LV, marked by dash lines) in the Mut brain. **b**, Serial coronal sections from WT, Het, and Mut mice. In the left half of each brain section, the brain slice area, the LV area, and the third ventricle area are highlighted by brown, blue, and red, respectively. **c**, The plot of the LV areas and the overall brain slice areas in serial coronal sections from 2M mice of different genotypes. Numbers in brackets show the numbers of animals analyzed for each genotype. **d**, Comparison of LV volumes between different genotypes. Each data point represents the result from one animal. **e**, Comparison of the volumes of the dorsal third ventricles (D3V) between different genotypes. **f**, Comparison of the LV volumes (mean ± s.e.m) between WT and Mut mice at different ages. **g**, Microscopic graphs of Nissl-stained coronal sections containing the aqueduct (Aq), D3V and the sub-commissural organ (SCO), and the fourth ventricle (4V). **h**, Comparison of the area of Aq, SCO, and 4V between WT and Mut mice. ns: not significant, Student *t*-test. **i**, The plot of the LV areas and the total brain slice areas of serial coronal sections of brains from WT and Mut female mice. **j**, Comparison of the volumes of LVs (left) and D3Vs (right) of WT and Mut female mice. Data are presented as mean ± s.e.m. * *p*<0.05, ** *P*<0.01, *** *P*<0.001, **** *P*<0.0001, ns: not significant; two-way ANOVA test in **c** and **i**, one-way ANOVA test in **d**, and Student *t*-test in **e**, **f**, **h**, **j**.

TUNEL staining and immunofluorescent staining of cleaved Caspase 3 did not show an increase in cell apoptosis in brains of Mut mice at P5 (Extended Data Fig. 4). Despite the expression of *Katnal2* in NPCs and OPCs during brain development, BrdU incorporation assays indicated normal progenitor proliferation in the VZ/SVZ of Mut mice at embryonic day 17 (E17) (Extended Data Fig. 5a-e). Moreover, there was no apparent difference in the density of PDGFR-α^+^ OPCs in the cortex, corpus collosum, and SVZ of Mut mice compared to WT littermates (Extended Data Fig. 5f, g). Together with the normal thickness and lamination of the cortex of Mut mice, these findings suggest that the hydrocephalus of *Katnal2*-deficient mice is not associated with abnormal neuro-gliogenesis during early development.

Although aqueduct stenosis is often associated with obstructive hydrocephalus^9^, we did not observe such stenosis in Mut mice (Fig. 4g). The subcommissural organ (SCO) is a circumventricular structure located anterior to the Sylvian aqueduct. EpCs of SCO secrete glycoproteins to facilitate CSF flow, and SCO dysplasia contributes to congenital hydrocephalus^28^. In Mut mice, the size of SCO didn’t differ from that of WT and Het littermates, but it appeared stretched in shape, likely due to ventricle dilation (Fig. 4g, h).

Excessive CSF secretion from the ChP is another potential cause of hydrocephalus^8, 29^. To investigate this, we examined the adherence junctions between ChP epithelial cells using immunofluorescent staining of β-catenin and zonula occludens 1 (ZO-1) and found that they were largely normal in both Mut mice and KO mice (Extended Data Fig. 6a). We also examined the polarized distribution of the basolateral Cl^-^-bicarbonate exchanger AE2 and the apical Na^+^ pump β1 subunit in the ChP, which plays a role in cross-epithelial CSF secretion. We found that their polarized distribution in the ChP epithelium was largely unaffected in Mut mice (Extended Data Fig. 6a). CSF is secreted under the osmotic gradient across the ChP epithelium. We measured the osmolality of CSF from 2-month-old mice and did not observe a significant difference between genotypes (Extended Data Fig. 6b, c). Therefore, the hydrocephalus of *Katnal2* deficient mice does not appear to be a result of abnormalities in the barrier structure of the ChP or the CSF secretion from ChP.

## Reduced CSF flow and ependymal ciliary motility of *Katnal2* mutant mice

Abnormal CSF circulation is known to cause ventricle dilation. To assess whether *Katnal2* mutation affects the bulk CSF flow, we injected 4 μl of Evans Blue (4 mg/mL) into the lateral ventricle (LV) of P7 mice and tracked the spread of the dye throughout the cerebral ventricles using serial coronal brain sections (Fig. 5a). In WT mice, the dye diffused rapidly and reached 4V within 2 min. However, in Mut mice, the Sylvian aqueduct and the 4V showed a weaker Evans Blue signal, while the LVs exhibited a higher signal compared to WT littermates (Fig. 5a). A similar reduction in Evens Blue diffusion was observed in *Katnal2* KO mice compared to WT littermates at P7 (Fig. 5a). These results suggest that the speed of CSF flow is reduced by *Katnal2* gene deficiency.

**Fig. 5:**
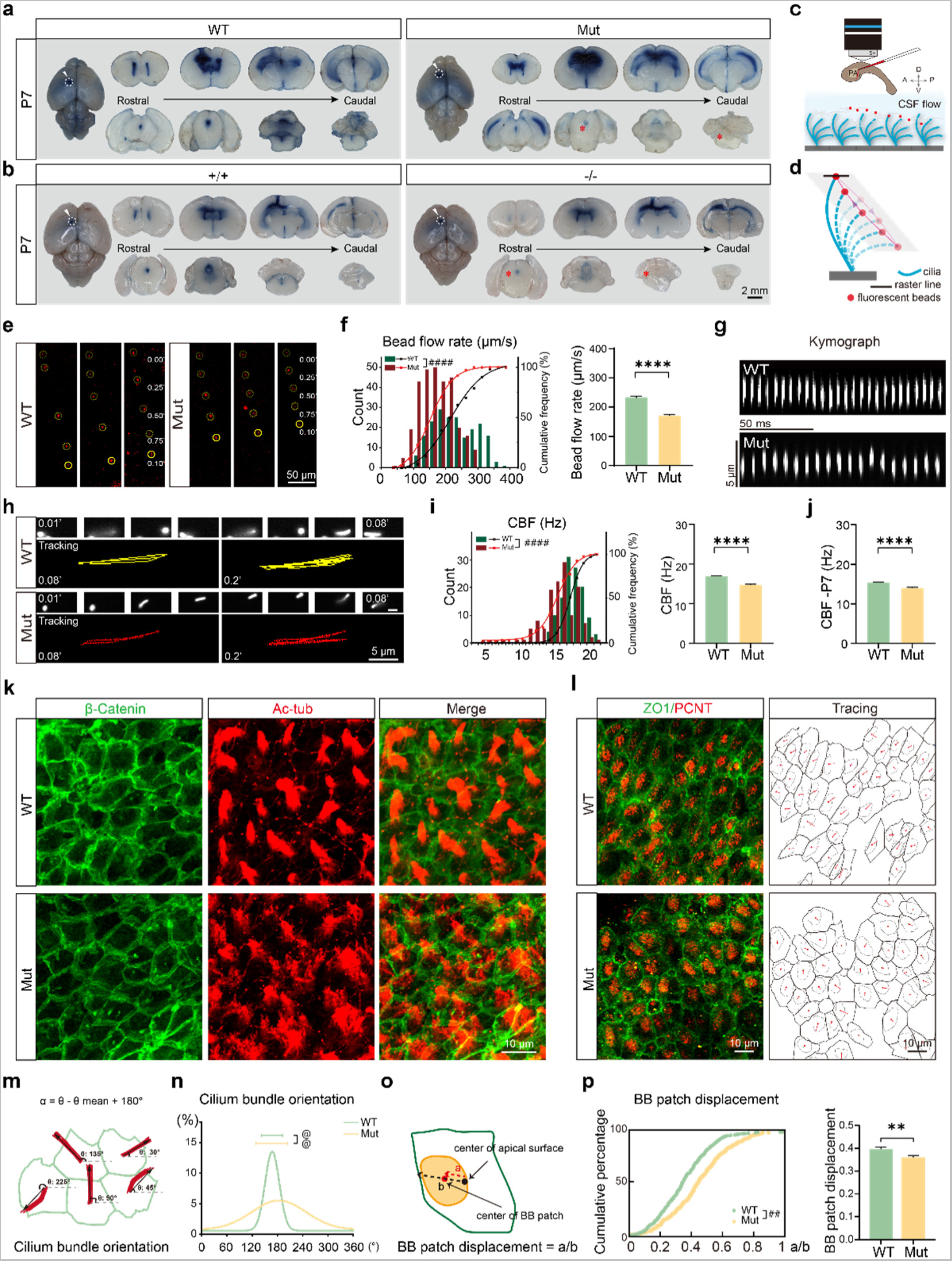
*Katnal2* mutation reduces CSF flow and ciliary motility and impairs the polarization and cilium organization of EpCs. **a**, Reduced Evans Blue (EB) diffusion in the ventricular system of *Katnal2* Mut and *KO* (-/-) mice compared to WT littermates at P7. Dashed line circles indicate the EB injection sites. Compared to WT littermates, *Katnal2* Mut and KO mice had less dye reaching the Aq and 4V regions (asterisks) 2 min after EB injection. Similar effect was observed in 3 out of 4 Mut/WT pairs and 2 out of 2 KO/WT pairs. **b**, Schematics of monitoring ependymal flow using fluorescent beads applied into a whole-mount preparation of the LV. A, anterior; D, dorsal; P, posterior; PA, point of adhesion; V, ventral. **c**, Representative time-lapse images showing sequential positions of beads (red dots, highlighted by yellow circles) on the surface of the ependymal layer of WT and Mut mice. The time points of each frame of image are shown at right (in seconds). **d**, Comparison of the distribution and the average value of the bead flow speed between WT and Mut mice at 2M. WT: 201 beads in 4 mice; Mut: 300 beads in 6 mice. **e**, Schematics of tracking cilium beating by cilium tip-attached fluorescent beads. The black line crossing the bead indicates the raster line for generating the kymograph. **f**, Representative kymographs delineating the to-and-fro motion of cilium tip of WT and Mut mice. **g**, Representative trajectories of cilium tip of WT and Mut mice within 0.2 second. **h**, Comparison of the distribution and average value of cilium beating frequency (CBF) between WT and Mut mice at 2M. WT: 120 cilia from 5 mice; Mut: 134 cilia from 6 mice. **i**, Comparison of CBF between WT and Mut mice at P7. WT: 218 cilia from 7 mice; Mut: 206 cilia from 7 mice. **j**, Immunofluorescent staining of cilium bundles of EpCs at rostral ventricle wall of WT and Mut mice. Red: acetylated tubulin (Ac-tub, show cilia); Green: β-catenin (show EpC shape). **k,** Schematics of the measurement of the cilium bundle orientation based on Ac-tub staining. **l**, Comparison of the distribution of cilium bundle orientations between WT and Mut mice. Error bars above the distribution curves show circular SD (CSD). WT: CSD = 50°, n = 576 cilium bundles from 5 mice; Mut: CSD = 75°, n = 713 bundles from 7 mice. **m**, Immunofluorescent staining and tracing of EpCs and their basal body (BB) patches. ZO1: tight junction protein 1, for EpCs; PCNT: pericentrin, for BBs. Red arrows indicate the BB patch displacement vector of EpCs. **n**, Schematics of the BB patch displacement vector (a) and the relative BB patch displacement (a/b). b: the cell radius in the direction of BB patch displacement. **o**, Cumulative distribution and mean value of the BB patch displacement of WT and Mut mice at 2M. WT: n = 341 cells from 5 mice; Mut: n = 396 cells from 4 mice. Data in histograms are mean ± s.e.m. **** *P* < 0.0001, Student’s *t* test. #### *P* < 0.0001, ## *P* < 0.01, Kolmogorov-Smirnov test. ^@@^ *P* < 0.01, Watson’s U2 two-sample test.

The beating of ependymal cilia creates regular and directed patterns of CSF flow near the ependymal surface. To investigate the ependymal flow in mice with *Katnal2* mutation, we tracked the motion of fluorescent microbeads applied to the surface of the ependymal layer in an acute whole-mount preparation of LVs^30^ (Fig. 5b). We observed an overall directional movement of fluorescent beads at the ependymal surface in Mut mice. However, the average velocity of bead translocation was significantly slower in Mut mice compared to WT littermates (Fig. 5c, d and Extended Data Video 1), indicating a reduced ependymal flow in Mut mice. These findings provide further evidence of impaired CSF dynamics in the absence of functional Katnal2.

In a brain slice preparation, high-speed time-lapse microscopic imaging of EpCs along the lateral wall of the LVs showed active beating of ependymal cilia in both WT and Mut mice. However, the cilium beating in Mut brain slices appeared to be less active compared to that of WT brain slices (Extended Data Video 2). To quantitatively analyze cilium beating, we tracked the motility of individual cilium of EpCs in the whole-mount preparations of LVs by monitoring the trajectory of microbeads attached to cilium tips (Fig. 5e-g)^31^. By examining kymographs and analyzing the to-and-fro motion of cilium tip-attached beads, we determined the frequency of cilium beating (CBF) (Fig. 5h and Extended Data Video 3). We found that CBF was significantly reduced in Mut mice compared to WT mice at both 2-month-old and at P7 (Fig. 5h, i). Collectively, these results demonstrated reduced ependymal ciliary beating and slower CSF flow in Mut mice, which could contribute to the CSF accumulation and ventricle dilation observed in Mut mice.

## *Katnal2* mutation affects ependymal ciliary organization

Defective differentiation and ciliogenesis of EpCs can lead to abnormal CSF flow and hydrocephalus^32, 33^. The ciliogenic forkhead transcription factor Foxj1 is crucial for the maturation and maintenance of EpCs^34^. To investigate the differentiation and ciliogenesis of EpCs in Mut mice, we labeled matured EpCs with Foxj1 antibody, stained cilium bundles with the antibody against acetylated tubulin (ace-tub) and marked cell boundaries with the antibody against β-catenin^35^ (Extended Data Fig. 7a, b and Fig. 5j). Lateral wall tissues of equivalent locations (rostral, central, and caudal) of LVs from WT and Mut mice were compared. We observed no significant differences in the density of EpCs or the percentage of Foxj1^+^ EpCs among total EpCs at the lateral wall of LVs between Mut and WT mice (Extended Data Fig. 7b-d). However, we did notice that the cilium bundles in Mut mice appeared more scattered compared to the well-aligned cilium bundle morphology in WT mice (Fig. 5j). Quantitative analysis of cilium bundle orientation revealed a significantly higher deviation in Mut mice (Fig. 5k, l), indicating less uniform orientation of cilium bundles. Using the fluorescence intensity of ace-tub to roughly estimate the cilium density, we found no significant difference in the average ciliary density per EpC or per unit area at the lateral wall of LVs between Mut and WT mice (Extended Data Fig. 7e, f). Immunofluorescent analysis of basal body (BB) patches of EpCs showed a significant reduction in the BB patch displacement from the cell center in Mut mice compared to WT mice (Fig. 5m-o), suggesting reduced translational polarity of EpCs in Mut mice. Scanning electron microscopy (SEM) further confirmed the more scattered ciliary orientation in Mut mice (Extended Data Fig. 7g). However, the gross cilium morphology, average cilium number per bundle, average cilium length appeared normal in Mut mice (Extended Data Fig. 7g-i). Transmission electron microscopy demonstrated a normal “9+2” organization of microtubule filaments inside ependymal cilia of Mut mice (Extended Data Fig. 7j). These results suggest that *Katnal2* mutation impaired the polarization of EpCs and their cilium bundle organization without a significant impact on the basic cilium structure. Finally, immunofluorescent staining of adenylate cyclase type III (AC3) showed a normal density and morphology of primary cilia in the cortex and hippocampus of Mut mice (Extended Data Fig. 8).

## EpC-specific ablation of *Katnal2* in neonatal mice causes ventricular dilation

To explore the causal relationship between ependymal Katnal2 and ventricular dilation, we performed conditional ablation of *Katnal2* specifically in the EpC lineage in neonatal mice. To do this, we developed mice with a conditional allele of *Katnal2*, featuring exons 2-12 flanked by loxP sequences, and crossed them with the *Foxj1^CreER^*mice to induce the ablation of floxed exons specifically in the EpC lineage upon tamoxifen treatment (Fig. 6a, b, h). The specificity and efficiency of tamoxifen-induced recombination by *Foxj1^CreER^* were evaluated using the Ai14 reporter line. Application of tamoxifen to the pregnant mice (75 mg/kg) for 5 consecutive days from P0 to P5 led to effective milk-mediated delivery of tamoxifen to pups, resulting in a gene recombination rate >80% in EpCs of P10 pups (Extended Data Fig. 9). Tamoxifen treatment induced a reduction in the body weight of pups, an effect independent of the genotype (Fig. 6e). We observed that conditional ablation of *Katnal2* in the EpC lineage from P0 caused a significant enlargement of LVs at P10 compared to control littermates without the *Foxj1^CreER^* allele (Fig. 6c, d), whereas the brain weight and cortex thickness were not affected (Fig. 6f, g). As a negative control, treatment with corn oil did not affect the ventricle size (Fig. 6c, d). Thus, ependyma-specific ablation of *Katnal2* is sufficient to drive hydrocephalus in neonatal mice. In 1-month-old mice with both Cre and homozygous conditional allele (*Foxj1^C^*^reER^::*Katnal2*^flox/flox^), tamoxifen injection (75 mg/kg for 5 consecutive days) to induce *Katnal2* ablation in the ependymal lineage failed to alter the ventricle size (Fig. 6h-j), suggesting that Katnal2 affects the CSF flow and ventricle size mainly during the early postnatal stage.

**Fig. 6.**
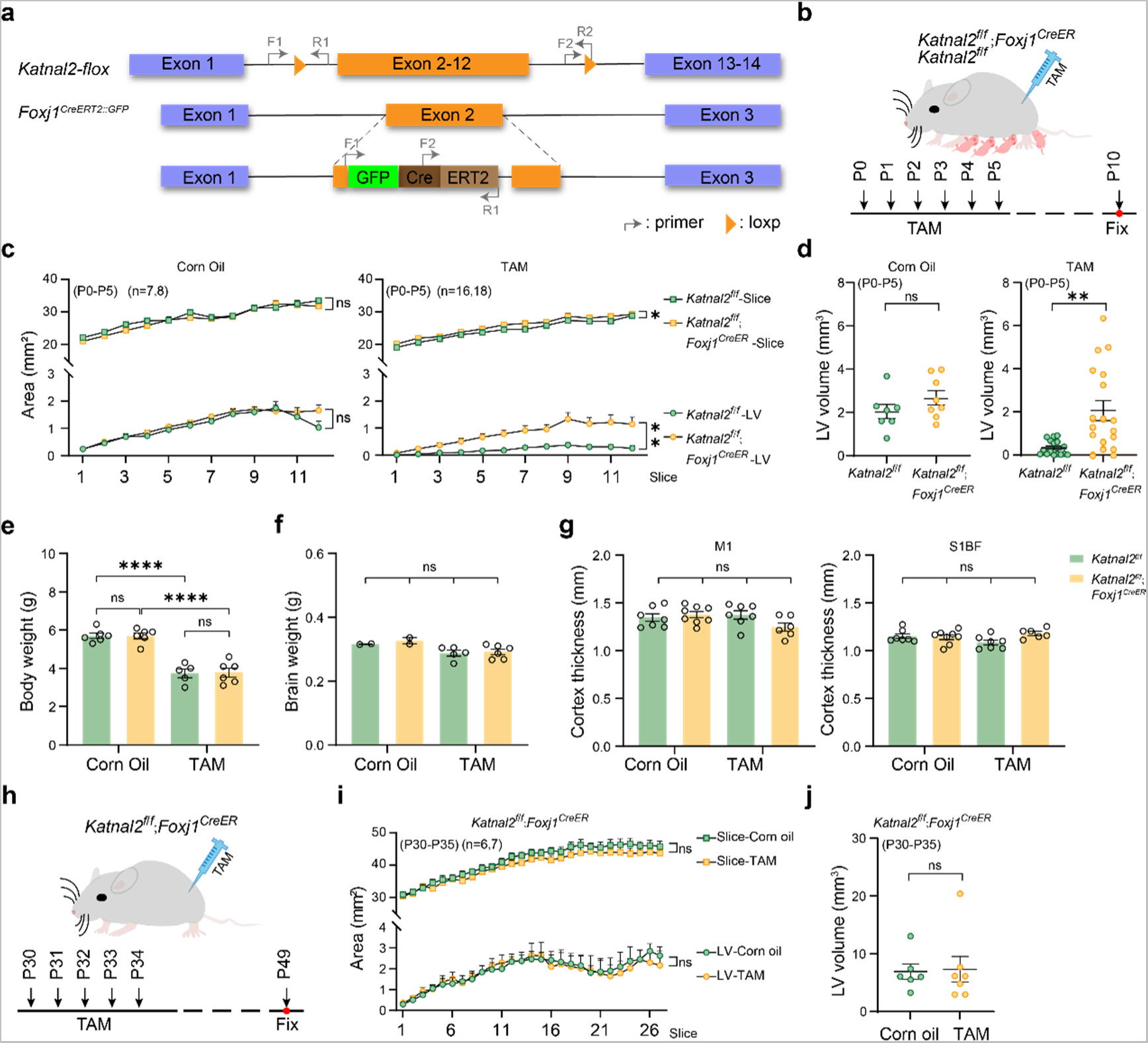
Ablation of *Katnal2* specifically in the ependymal lineage during the early postnatal stage results in brain ventricle enlargement. **a**, Schematics of the conditional allele (*Katnal2-flox*) and the Cre allele (*Foxj1^CreERT^*^2^*^::GFP^*). **b**, Schematics of the milk-mediated transmission of tamoxifen (TAM) to newborn mice of two different genotypes. **c**, Comparison of the LV areas and the overall brain slice areas in serial brain sections from mice of two genotypes with or without TAM treatment. Corn oil treatment as the negative control. **d**, Comparison of the LV volumes in P10 mice of two genotypes with or without maternal application of TAM. Each data point represents the result from one animal. **e-g**, Effects of tamoxifen treatment (P0-P5) on the body weight (**e**), brain weight (**f**), and cortical thickness (**g**) of P10 pups of different genotypes. **h**, Schematics of TAM treatment to induce conditional KO (CKO) in adult mice (*Foxj1*^CreER^::*Katnal2*^flox/flox^). **i**, Comparison of the LV areas and the overall brain slice areas in serial brain sections from adult CKO mice (TAM) with control mice (Corn oil). **j**, Comparison of the LV volumes in adult CKO mice with control mice. Data are presented as mean ± s.e.m. * *P*<0.05; ** *P*<0.01; **** *P*<0.0001; ns: not significant; two-way ANOVA test in **c** and **i**, Student *t*-test in **d-g** and **j**.

## Discussion

While most studies focus on neuronal deficits in ASD, the present study sheds light on the significant role of ependymal glia in the etiology of ASD. Our study revealed that mice with an ASD-associated mutation of *Katnal2* developed postnatal hydrocephalus, which can be attributed to impaired ciliary beating of EpCs at early postnatal stage. The observed hydrocephalus in *Katnal2* mutant mice aligns with the ventriculomegaly observed in some patients carrying genetic variants of the *KATNAL2* gene, including an autism-diagnosed patient^27^. These findings support the potential involvement of hydrocephalus, caused by the risk gene, in the pathogenesis of ASD. While a causal relationship between hydrocephalus and ASD has not yet been established, emerging evidence suggests such a possibility. Large-scale comparisons of anatomical MRI scans have revealed significantly larger ventricular volumes in individuals with ASD^4^. The ventricle enlargement has been confirmed by the ENIGMA ASD Working Group^36^ and longitudinal brain imaging studies across various age groups^5, 37^. Furthermore, larger brain ventricles have been identified in some infants and toddlers before an ASD diagnosis^6^. A recent population-based cohort study in Denmark reported a significant association between hydrocephalus and ASD, indicating a potential causal or shared etiology^7^. It’s important to note that not all ASD patients are diagnosed with ventriculomegaly, and the incidence of hydrocephalus in ASD patients may have been underestimated due to individual variation in ventricle volume, particularly in cases of mild to moderate hydrocephalus. Macrocephaly is frequently observed in children with ASD. While abnormal neurogenesis is often considered to underlie brain overgrowth and macrocephaly in ASD^38^, hydrocephalus could be another potential factor contributing to macrocephaly in some autistic children.

Recent studies have suggested that risk genes for congenital hydrocephalus converge on the proliferation and neurogenesis of NPCs in the embryonic cortex^11, 12^. However, since EpCs are also derived from embryonic NPCs, it is plausible that some risk genes of congenital hydrocephalus may impair the genesis of EpCs to cause the hydrocephalus. Previous studies have highlighted the crucial role of NG2^+^PDGFR-α^+^ OPCs in hydrocephalus^39^. In our study, we discovered that *Katnal2* is expressed not only in EpCs but also in NPCs and NG2^+^PDGFR-α^+^ OPCs. However, in *Katnal2* mutant mice, we did not observe a significant reduction in the density of BrdU^+^ cycling cells and PDGFR-α^+^ OPCs, nor did we observe increased cell apoptosis in mutant brains. The significant ventricular enlargement observed in the mutant mice begins during the first postnatal week, coinciding with the maturation of ependymal cilia at the lateral wall of LVs during normal development^40^. Finally, conditional ablation of *Katanl2* specifically in the EpC lineage from an early postnatal stage was sufficient to induce ventricular enlargement. These findings collectively suggest that the hydrocephalus observed in *Katnal2* mutant mice primarily stems from impaired ciliary beating and reduced CSF flow, rather than defects in the proliferation and neurogenesis of progenitor cells, although a modulatory role for Katnal2 in neuro- and glio-genesis cannot be entirely ruled out.

Several studies have established an association between various cilia genes and developmental brain disorders, including ASD^41, 42^. However, most of these studies have predominantly focused on the role of primary cilia, while the contribution of motile cilia in ASD etiology has been largely overlooked. The present study highlights the critical role of the high-risk ASD gene *Katnal2* in the polarized organization and beating of ependymal ciliary bundles. The effect of Katnal2 on ciliary organization and motility could be attributed to its involvement in establishing the planar cell polarity (PCP) of EpCs, which is crucial for the correct positioning of motile cilia^43^. A recent study demonstrated a delay in the convergent extension of the neuroepithelium during neurulation in *Katnal2* KO zebrafish^23^. Considering the well-established association of PCP genes with the convergent extension, this finding in Zebrafish implies a potential role of Katnal2 in the PCP pathway. Katnal2 belongs to the katanin family of microtubule-severing proteins; however, the predicted microtubule-severing activity of this protein has yet to be validated. The mechanism by which Katnal2 regulates EpC polarity and PCP pathway remains to be investigated.

While gene co-expression analysis suggests an important role of KATNAL2 in cilium assembly, we did not observe a prominent defect in the morphology and axoneme structure of ependymal cilia, unlike in sperm, where *Katnal2* is more abundantly expressed, and its mutation leads to severe deficits in flagellar morphology and motility. The absence of apparent morphological defects in individual ependymal cilia may be attributed to the low level of *Katnal2* expression in EpCs. Notably, defects in the ciliary structure are prominent in several mouse models of ciliopathy, which display severe brain ventricle swelling^44^. In contrast, *Katnal2*-deficient mice exhibit moderate ventricular dilation, indicative of a relatively milder impairment in ciliary function. In human patients, severe hydrocephalus during development may lead to more profound structural and functional deficits of the brain, resulting in more severe physical, psychological, and cognitive symptoms, rather than ASD. However, a sustained yet modest hydrocephalus might be more relevant to the ASD pathology, where the structural alteration of the brain is less prominent, and essential brain functions are largely maintained.

In summary, our study has revealed the significant role of the ASD risk gene *Katnal2* in brain development, particularly in its influence on ependymal ciliary organization and motility during the early postnatal stage. These findings indicate a potential link between ciliary defects, hydrocephalus, and the development of ASD. Further research is required to uncover the molecular mechanisms that govern the regulation of EpC polarity by Katnal2, as well as to explore potential therapeutic approaches aimed at targeting ependymal ciliary function for the treatment of ASD and other neurodevelopmental disorders.

## Methods

### Animals

All animal procedures were performed following guidelines for the Care and Use of Laboratory Animals of the National Institutes of Health and were approved by the Institutional Animal Care and Usage Committee of the East China Normal University (m20221003). Mice were maintained on a C57BL/6J background or on a C57BL/6J -CD-1 mixed background. Mice were raised in a facility at 23°C and 50% humidity with a 12 h/12 h light/dark cycle and access to food and water *ad libitum*.

Knock in of the *Katnal2* splice site mutation in C57BL/6J mice was conducted using the CRISPR/Cas9 genome editing technology. Briefly, gRNA (CCGGgtaagatctgctattcaat), single-strand donor DNA (TGAGTGCATTTATTGGCATGAACAGTGAGATGCGAGAACTGGCAGCGGTGGTGAGCCGGataagatctgctattcaattcacaaatttatggaggcagaccagggctcctggagcattgt), and Cas9 mRNA were introduced into the cytoplasm of mouse zygotes through microinjection using the Eppendorf TransferMan NK2 micromanipulator. Injected zygotes were transferred into pseudopregnant female mice after 2 h culture in KSOM medium. The founder mice were backcrossed to WT C57BL/6J mice for at least 5 generations to eliminate potential off-target mutations before experiments.

The *Katnal2*-null allele and the *Katnal2*-floxed allele were generated by the GemPharmatech Co., Ltd. (Nangjing, China). Briefly, to generate the *Katnal2*-null allele, two pairs of gRNAs (5s1: GCGTTTACCAGCTGCTATAC, 3s1: AGCCCGCCCTGAGGCTTCAT) were co-injected with Cas9 mRNA into mouse zygotes to induce the ablation of exons 2-12. To generate the *Katnal2*-floxed allele, Cas9 mRNA, two pairs of gRNAs (5s1: GCGTTTACCAGCTGCTATACAGG, 3s1: AGCCCGCCCTGAGGCTTCAT), and two single-stranded oligodeoxynucleotides (ssODNs) containing loxP sequences were co-injected into mouse zygotes to insert a pair of loxP sequences flanking the exon 2-12 region.

*Katnal2* conditional knockout mice were generated by crossing *Katnal2*-floxed mice with *Foxj1^tm^*^1.1^(cre/ERT2/GFP)*^Htg^*mice (Jackson #027012, termed as *Foxj1^CreER^* in this study). *Ai14* reporter line (Jackson #007914) was crossed with the *Foxj1^CreER^*line to report the Cre recombinase activity. To induce Cre activity in newborn mice, 75 mg/kg of tamoxifen (10 mg/ml in corn oil) was injected subcutaneously into the mother mice daily for 5 consecutive days to be transmitted to pups through the milk. To induce Cre activity in adult mice, daily injection of 75 mg/kg of tamoxifen was applied subcutaneously for 5 consecutive days.

For genotyping, genomic DNA was extracted from mouse tail clips after proteinase K digestion. For the *Katnal2* splice site mutation allele, PCR amplification of genomic DNA was conducted with the primer pair: 5′-CCAGGCACCATTTGTTTCG-3′ and 5′-CCTGCACGCTTTGCTTTTC-3′, followed by sequencing of the PCR product. Primers for the genotyping of the *Katnal2*-null colony are F1 (5′-ACGCCTAGCAGTGTGAATGAG-3′) and R1 (5′-TAGAGAGGTTCTCCTTCCTGCTG-3′) for the WT allele and F2 (5′-TCAGGGCGGGCTCCTTCTG-3′) and R1 for the mutant (Mut) allele. The sizes of PCR products for the WT and Mut alleles were 132 bp and 283 bp, respectively. The primers for genotyping of the *Katnal2*-floxed colony are F1 (5′-TATCTGGAGCTGTCTTCGAGCAC-3′) and R1 (5′-AGACAGCCTCATTCACACTGCTAG-3′) for the WT allele and F2 (5′- GCATCGCATTGTCTGAGTAGGTG-3′) and R2 (5′-ACTATGTAGACCAGCCTGGCATC-3′) for the floxed allele. The sizes of PCR products for the WT and floxed alleles were 312 bp and 417 bp, respectively. The primers for genotyping of the *Foxj1^CreER^* mice are F1 (5′-GCAGATGGAGAGAGGTGGAG-3′) and R1 (5′-CTTGGCGTTGAGAATGGAGA-3′) for the WT allele and F2 (5′-ATTGCATCGCATTGTCTGAG-3′) and R1 for the mutant allele. The sizes of PCR products for the WT and Cre alleles were 472 bp and 694 bp, respectively. Primers for genotyping of the Ai14 reporter line are F1 (5′-AAGGGAGCTGCAGTGGAGTA-3′) and R1 (5′-CCGAAAATCTGTGGGAAGTC-3′) for the WT allele and F2 (5′-CTGTTCCTGTACGGCATGG-3′) and R2 (5′-GGCATTAAAGCAGCGTATCC-3′) for the reporter allele. The sizes of PCR products for the WT and reporter alleles were 297 bp and 196 bp, respectively.

### RT-PCR

Total RNA was extracted from whole brains of mice using Trizol agent (Takara, Japan) and treated with DNase following the manufacturer’s instructions. For cDNA synthesis, the FastQuant RT Kit (Tiangen Biotech, Beijing, China) was used. RT-PCR and cDNA sequencing were performed following standard methods. Quantitative real-time PCR (qPCR) was performed with the CFX Opus 96 Real-Time PCR System (Bio-Rad) using the FastKing One Step RT-qPCR kit (SYBR Green, Tiangen Biotech). Primers for qPCR are *Katnal2*: F 5′-CTTTGAATGTAACCCAGACC-3′, R: 5′-CACAACGGCTTCTTTACT-3′; *Gapdh*: F: 5′-GTGGAGTCATACTGGAACATGTAG-3′, R: 5′-AATGGTGAAGGTCGGTGTG-3′.

### RNAscope *in-situ* hybridization

For the detection of *Katnal2* mRNA, two different kits were used: the RNAscope 2.5 HD Brown Reagent Kit (#322300, Advanced Cell Diagnostics, USA) for chromogenic detection and the RNAscope Multiplex Fluorescent V2 Assay kit (#323100, Advanced Cell Diagnostics) for fluorescent detection. Paraffin sections, with a thickness of 5 μm, ranging from E13.5 to adult stages, were mounted on Superfrost-plus adhesion glass slides (Fisher, USA). The detection of *Katnal2* mRNA was performed following the manufacturer’s protocols for the respective kits. Mouse-specific *Katnal2* probe was designed and synthesized by the manufacturer. Positive (mouse *Ppib*) and negative (*DapB*) control probes were included in each experiment. The procedure started with the baking of the sections at 60°C for 1 h, followed by deparaffinization. Hydrogen peroxide treatment was carried out at RT for 10 min. Target retrieval was performed by heating the sections at 100°C for 15 min, followed by protease treatment at 40°C for 15 min. Probes were then hybridized with the sections at 40°C for 2 h, followed by six steps of amplification. For chromogenic signal detection, the sections were subjected to 3,3’-diaminobenzidine (DAB) incubation. After washing in water and drying, the slides were mounted with Ecomount (Biocare Medical, Concord, CA, USA) and stored at RT. For fluorescent signal detection, the sections were incubated with TSA® Plus Cy5.

### Gene co-expression analysis

The gene co-expression analysis was conducted using the BrainSpan human brain transcriptome dataset (RNA-Seq Gencode v10) available at www.brainspan.org. This dataset comprises 256 transcriptomes obtained from 16 different brain regions, covering a wide developmental range from post-conception week 8 to 40 years old. Gene expression levels were quantified using RPKM (reads per kilobase per million mapped reads). To ensure robust analysis, low abundance genes with an average expression across different tissue samples of less than 1 RPKM were filtered out. The remaining dataset included a total of 12,250 genes for further analysis. The Pearson’s correlation coefficient of each gene in the entire genome with *KATNAL2* was calculated based on the gene expression levels in each brain sample. The top 1000 genes with the highest correlation coefficients were selected for Gene Ontology (GO) analysis using DAVID v6.8 (http://david.ncifcrf.gov/tools.jsp). The human whole-genome genes provided by DAVID were used as background genes for the GO analysis. A Benjamini-Hochberg corrected p-value of 0.05 was used as the threshold for significance in the GO analysis.

### Single-cell RNAseq

Single-cell RNA sequencing (scRNAseq) was conducted using the 10x Genomics platform according to the manufacturer’s instructions. Briefly, P5 cortical tissues were treated first by 0.25% Trypsin-EDTA at 37℃ for 30 min, then by 0.2% type-II collagenase for 4 h to dissociate the cells. After removing cell clumps using a 70 m sterile nylon mesh cell strainer, single cells were isolated using fluorescence-activated cell sorting and encapsulated in droplets along with barcoded beads and reverse transcription reagents using the 10x Genomics Chromium Controller. RT-PCR were performed within the droplets, followed by fragmentation and library preparation using the 10x Genomics Single Cell 3’ Reagent Kit. Sequencing was performed on an NovaSeq 6000 Sequencer (Illumina San Diego, California, USA) with a read length of 150 bp. Raw reads were processed using the Cell Ranger software (v6.1.1) to generate gene-barcode matrices for each individual cell. Cells with low quality (>30% mitochondrial or ribosomal content) or low gene count (<200 unique genes) were removed from downstream analysis. Gene expression was normalized and log-transformed using the Seurat package (version 3.2.2) in R software (v4.0.2). Genes that were expressed in fewer than 3 cells were excluded from further analysis. After principal component analysis (PCA). Differential gene expression analysis was performed using the Wilcoxon rank sum test implemented in the Seurat package. Markers of each cluster were identified using the FindAllMarkers function based on the fold change (>2) and adjusted p-value (<0.05). Tailored cell type markers (Extended Table 3) were used for annotation of each cell cluster.

### Testis histology and sperm analysis

Testes were collected from adult male mice following euthanasia and immediately fixed in 4% paraformaldehyde (PFA) at 4°C overnight. The fixed tissues were dehydrated using a 30% sucrose solution and then embedded in O.C.T. compound (Sakura, Japan). Subsequently, the testes were sectioned into 10 μm thick slices using a cryostat microtome (CM1850, Leica). The obtained slices were carefully mounted onto glass slides and stored at -20°C until further analysis. For histology analyses, the slices were subjected to routine processing for hematoxylin and eosin (H&E) staining.

The cauda epididymis was isolated from adult male mice immediately following euthanasia. It was then cut into small pieces to facilitate the release of spermatozoa, which were allowed to swim out in a 24-well plate containing approximately 0.5 ml of warmed Tyrodes’ solution (Sigma, USA) per well. The plate was placed in an incubator with 5% CO2 at 37°C for 15 min to promote sperm motility. After incubation, the supernatant with no apparent tissue debris was carefully transferred to 1.5 ml Eppendorf tubes. For Giemsa staining, 10 μl of the supernatant containing spermatozoa was mixed with Giemsa dye solution (Beyotime, China) and incubated for 10 min at RT. The mixture was then spread onto a glass slide and allowed to air-dry at RT. For morphology analysis, at least 30 spermatozoa from each sample were examined using standard criteria.

For the analysis of sperm motility, the obtained supernatant containing spermatozoa was transfered into a specialized chamber with a depth of 20 μm (Leja, 025107-025108, IMV Technologies, USA). The chamber allows for the observation of sperm movement in a controlled environment. The movement of sperm heads was tracked using the IVOS sperm analysis system (Hamilton Thorne, USA) at a rate of 100 fps. To ensure robust statistical analysis, a minimum of 300 spermatozoa from six objective fields within each chamber were recorded to accurately capture the motility patterns. The major sperm motility parameters recorded for further analysis included total motility (%), progressive motility (%), average path velocity (VAP, μm/s), straight line velocity (VSL, μm/s), and curvilinear velocity (VCL, μm/s).

### Brain histology and immunofluorescence

Anesthetized mice were subjected to perfusion with PBS, followed by 4% PFA. The brains were then cryoprotected in a solution of PBS containing 30% sucrose and subsequently embedded in O.C.T. compound for sectioning. Sections of 30 μm thickness were obtained using a cryostat microtome (CM1850, Leica) for Nissl staining and immunofluorescence microscopy. For antigen retrieval, brain sections were treated with citrate buffer at 95-100°C for 20 min. The sections were rinsed with PBS and permeabilized with 0.3% Triton X-100 for 30 min at RT. Subsequently, the sections were blocked with 10% goat serum and incubated with primary antibodies at an appropriate concentration at 4°C for 48 h. Following thorough rinsing with PBS, sections were incubated with Alexa Fluor 488/546-conjugated secondary antibodies for 2 h at RT. After a thorough wash with PBS, the brain sections were mounted for confocal imaging with the Olympus FV10i system. The antibodies used for immunofluorescence are summarized in Extended Data Table 4.

### CSF collection and analysis

Adult mice were anesthetized and secured onto an automated stereotaxic instrument (#71000, RWD, China) with the use of ear bars to stabilize the head. The head position was adjusted to orient the basilar region upwards. A scissor was used to make an incision in the skin and upper muscle layer, revealing the skull base covered by a thin layer of muscle. The thin muscle layer was carefully removed using fine forceps under a stereoscope, ensuring minimal bleeding, to expose the cisterna magna (CM). Using the stereotaxic instrument, a sharp glass micropipette with a tip diameter of 20 μm was inserted into the CM. Approximately 10-15 μl of clear CSF was collected from each animal using a 1 ml syringe connected with the micropipette. The collected CSF samples were frozen at -80 °C until analysis. Osmolality measurements were performed using an automated freezing point osmometer (OsmoTECH XT, Advanced Instruments, USA).

### Systemic CSF flow analysis

Mice were deeply anesthetized with intraperitoneal administration of sodium pentobarbital (40 mg/kg). Evans Blue (4 mg/mL) was injected into the left lateral ventricle at a rate of 4 μl/min. The injection coordinates for P7 mice were -2.0 mm depth, -0.8 mm left, and -0.1 mm posterior from the Bregma. Mice were promptly decapitated 2 min after injection. The brains were fixed in 4% PFA overnight and then cut into 2 mm thick coronal slices using a brain matrix slicer. The brain sections were mounted for imaging using a stereoscope (M165FC, Leica).

### Fluorescent microbeads-based assays of the ependymal flow

Whole-mount brain ventricle preparations were utilized to observe the dynamic CSF flow on the surface of the ependymal layer as described previously^30^. The hemisphere containing the lateral ventricle (LV) was carefully dissected from anesthetized mice and pinned in a dish with L-15 culture media at RT. Fluorescent latex beads (1-μm diameter, Sigma L2778) were gently released onto the surface of the LV wall using a syringe pump controller (TJ-4A, LongerPump, Hebei, China). The movements of beads were monitored using a 60x water immersion objective on a fluorescence microscope (BX51WI, Olympus) and recorded using a CMOS camera (optiMOS^TM^-RM16C, Teledyne Qimaging, Tucson, AZ) at a rate of 100 frames per second (fps).

### Monitoring ciliary beating

To monitor ependymal ciliary beating, the mouse brains were rapidly removed following anesthesia and decapitation of the animal. Coronal brain slices were obtained at a thickness of 300 μm using a vibratome (VT 1200S, Leica) in ice-cold oxygenated cutting solution (228 mM sucrose, 11 mM glucose, 26 mM NaHCO_3_, 1 mM NaH_2_PO_4_, 2.5 mM KCl, 7 mM MgSO_4_, and 0.5 mM CaCl_2_) with a continuous supply of 95% O_2_ and 5% CO_2_. The obtained slices were then allowed to recover for over 30 min in oxygenated artificial cerebrospinal fluid (ACSF: 119 mM NaCl, 2.5 mM KCl, 1 mM NaH_2_PO_4_, 1.3 mM MgSO_4_, 26 mM NaHCO_3_, 10 mM glucose, and 2.5 mM CaCl_2_) at 28°C before imaging. Subsequently, the brain slices were transferred to a perfusion chamber mounted on a microscope (BX51WI, Olympus). Phase-contrast imaging of the lateral ventricle (LV) surface was performed using a 60x water immersion objective. High-speed videography was carried out using a CMOS camera (Teledyne Qimaging) at an acquisition rate of 100 fps.

To analyze the beating pattern of individual cilium, fluorescent latex beads were diluted at a 1:200 ratio with L-15 medium and applied to the whole-mount sample of the LV. After waiting for 5 min, the fluorescent microbeads settled and attached to the tips of the cilia. Subsequently, the LV walls were pinned in a dish containing fresh medium at RT. The cilium tip movements were captured using the above-described high-speed videography setup. The resulting video was converted to an image stack using Image J software. To graphically illustrate the back-and-forth motion of the beads attached to the cilium tips, kymographs were generated by drawing a scan line through the selected fluorescent beads in the image stack.

### Electron microscopy

For scanning electron microscopy (SEM), the ependyma lining of the LV was dissected and fixed in 2% PFA (electron microscopy grade, Sigma) and 2.5% glutaraldehyde in 0.1 M PBS (pH 7.4) at 4°C overnight. Tissues were washed thoroughly in PBS, then post fixed in 0.5% osmium oxide (in 0.1 M PBS) for 1 h. Samples were washed thoroughly in PBS and dehydrated prior to critical point drying in 100% ethanol. Tissue blocks were gold palladium coated using a sputter coater and scanned with a scanning electron microscope (S4800, Leica).

For transmission electron microscopy (TEM), samples were further embedded in Epon 812 embedding resin (Electron Microscopy Sciences, USA). Ultra-thin sections were obtained using a LEICA UC6 ultramicrotome. Sections were placed on 200 mesh grids and stained with uranyl acetate and lead citrate, then imaged using a transmission electron microscope (JEM2100, Jeol, Japan).

### Antibodies and reagents

Primary and secondary antibodies used in this study are summarized in Extended Data Table 4. In addition, the following reagents were used: BrdU (Abcam, ab125467) for cell proliferation assay, Evans Blue (Sangon Biotech, Shanghai, A602025-0005) as indicative dye, and tamoxifen (Sigma, T5648) for Cre-recombination induction.

### Behavioral analysis

Behavioral tests were conducted on mice aged 2-3 months. To minimize anxiety levels, mice were handled for 10 min daily over a period of 3 days prior to the tests. The tests were performed in the following order: open field test, elevated plus maze, sociability test, rotarod test, balance beam test, gripping strength test, and resident-intruder test. After completing each test, animals were given a minimum of 3 days of rest. The behavioral apparatus was disinfected with 70% ethanol prior to the experiments and cleaned with 10% ethanol between tests. A waiting period of at least 5 min was observed to allow the evaporation of ethanol before commencing the next test. Throughout the experiments, the experimenter remained blind to the genotype or treatment of the mice.

#### Open field test (OFT)

OFT was performed in open field arenas made of Plexiglas (25 cm × 25 cm × 30 cm), and mouse behavior was tracked by the ActiTrack software (Panlab, Spain). This experimental system can record mouse behaviors in 4 arenas simultaneously. For each animal, the total distance moved, the time and locomotion distance in the center of the arena (1/2 of total size) within 30 min of free exploration of the arena were recorded.

#### Elevated plus maze (EPM)

A standard EPM apparatus was used for the test. The maze was 30 cm above the floor, consisting of two open and two closed arms, 30 cm × 5 cm each, connected by a 5 cm × 5 cm central platform. The test mouse was gently placed on the central platform with its head facing one closed arm and was allowed to freely explore for 5 min. Mouse behaviors were tracked by the Any-Maze system (Stoelting, USA). The time of open arm exploration for each animal was recorded.

#### Sociability test

The standard 3-chamber sociability test was performed using a 60 cm × 30 cm × 40 cm white Plexiglas arena divided into three chambers (20 cm × 30 cm × 40 cm) by clear Plexiglas dividers with an open door connecting each chamber. The test mouse was first habituated to the middle empty chamber for 5 min. In the first stage, there was only an empty metal wire cage in each side chamber. The test mouse was allowed to explore freely for 10 min. At the second stage, a same-sex stranger mouse was placed under the cage of one side chamber, permitting the test mouse to freely explore for 10 min. At the third stage, another same-sex stranger mouse was placed under the cage of the other side chamber, allowing the test mouse to freely explore for another 10 min. Mouse behaviors were recorded with an overhead camera and tracked using Any-Maze software. We recorded the total time that the test mouse spent exploring each side chamber and the total time sniffing stranger mice or empty cages for further statistical analysis.

#### Rotarod

The rotarod test was conducted using a rotarod apparatus (JLBehv-RRTG, Shanghai Jiliang Software Technology, China). The rotation of the rod was set to accelerate from 4 RPM to 40 RPM over the course of 300 seconds. All mice were tested on the accelerating rod five times per day for 7 consecutive days. For each animal, the latency for keeping balance on the rod in each test was recorded.

#### Grip Strength Test

The test mouse was placed on a metal wire mesh. The mesh was quickly inverted, and the latency for the animal to hang to the mesh was recorded. The maximum hanging time was set to 3 min, after which the trial was stopped. For each animal, 5 trials were carried out, and the average hanging time was calculated.

#### Marble burying

Standard polycarbonate rat cages (26 cm x 48 cm x 20 cm) with fitted filter-top covers were used for the marble burying test. Add fresh, unscented mouse bedding material to each cage to a depth of 5 cm. Place glass toy marbles (assorted styles and colors, 14 mm diameter, 5.2 g in weight) gently on the surface of the bedding in 5 rows of 4 marbles. Place one mouse in a corner of the cage containing marbles. Allow the mouse to remain in the cage undisturbed for 30 min. Remove the mouse and count the number of marbles buried. Score a marble as buried if two-thirds of its surface area was covered by bedding.

## Statistical analysis

Quantitative data analyses were performed by an experimenter who was blind to the experimental conditions whenever feasible. Statistical analyses were conducted using Prism 8.0 (GraphPad Software Inc, USA). For all statistical analysis, two-tailed tests were used. One-way ANOVA with Tukey’s post-hoc test was employed for comparisons among multiple groups. Two-way ANOVA was employed to analyze the group difference in serial coronal brain sections. Normal distribution of data was validated. Statistical significance was set at *P* < 0.05. For representative images, similar results were obtained in at least two independent trials. Detailed information of statistical analysis for each experiment is shown in figure legends and in the Supplementary Information.

## Acknowledgments

This work was funded by the National Key Research and Development Program of China (2022YFC2705200) and the National Science Foundation of China (31871501, 32061143016). We thank ECNU Multifunctional Platform for Innovation (010 and 011) for the mouse transgenic and breeding service.

## Author contributions

S. Qiu, H. Cao, Y. Xue, N. Wu, S. Xie, M, Song and J. Ma conducted experiments; S. Qiu and Y. Xue conducted data analysis; S. Qiu and Y. Pan contributed to figure organization; S. Qiu, J. Ma, and X. Yuan contributed to the writing of the manuscript; X. Yuan and P. Baas conceived the project; X. Yuan and J. Ma supervised the project.

## Additional information

Supplementary Information is available for this paper. Correspondence and requests for materials should be addressed to Dr. Xiao-Bing Yuan

**Data availability**: All raw data for Figs. 1-6 and Extended data Figs. 1-9 are provided in Extended data Tables 1-3 and Supplementary Information. Antibodies used for immunofluorescent staining in this study were provided in Extended data Table 4.

## Extended data figure/table/movie legends

**Extended data Fig 1.**
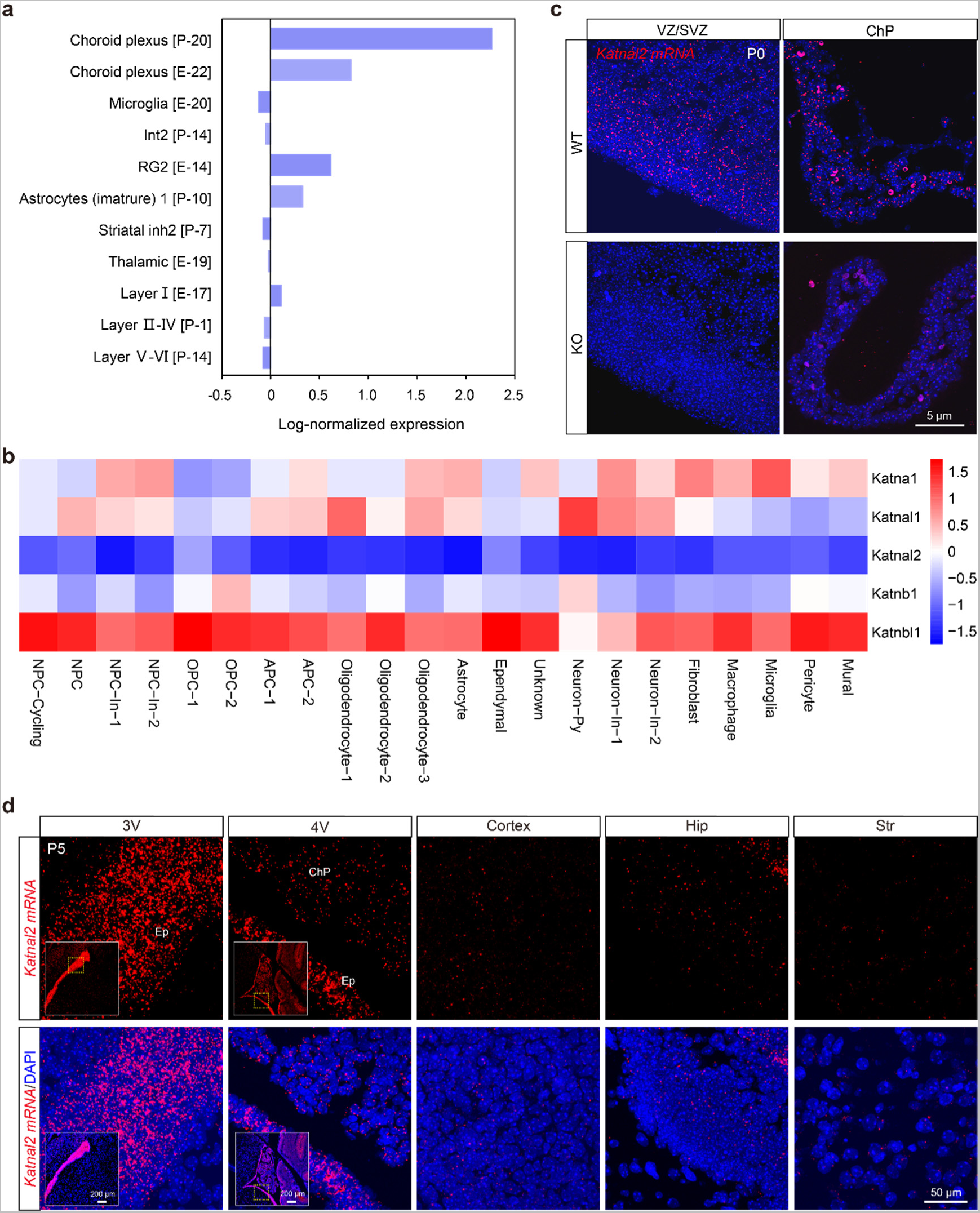
Single-cell transcriptomic analysis and *in-situ* hybridization of *Katnal2* in mouse cortex. **a**, Single-cell transcriptomic analysis of *Katnal2* in mouse brains at E14.5 and P0. Data are sourced from Zylka lab (http://zylkalab.org/datamousecortex)^25^. **b**, Heat-map presentation of relative expression levels of different *Katanin* family members in each cell cluster of mouse cortex at postnatal day 5 (P5) based on scRNA-seq. In each cell cluster the normalized expression level (Z-score) of each gene was scaled across *Katanin* family members and coded by pseudo color. **c**, Fluorescent *in-situ* hybridization of *Katnal2* (red) in the VZ/SVZ and choroid plexus of wildtype (WT) and *Katnal2* Knockout (KO) mice at P0. VZ/SVZ, ventricular and subventricular zone; ChP, choroid plexus. **d**, Fluorescent *in-situ* hybridization of *Katnal2* (red) in WT and KO mouse brains at P5. High-magnification images of the 3V and 4V regions are from the selected areas of low-magnification images of the brain slice shown in the inserts. 3V, third ventricle; 4V, fourth ventricle; Hip, Hippocampus; Str, striatum.

**Extended Data Fig 2.**
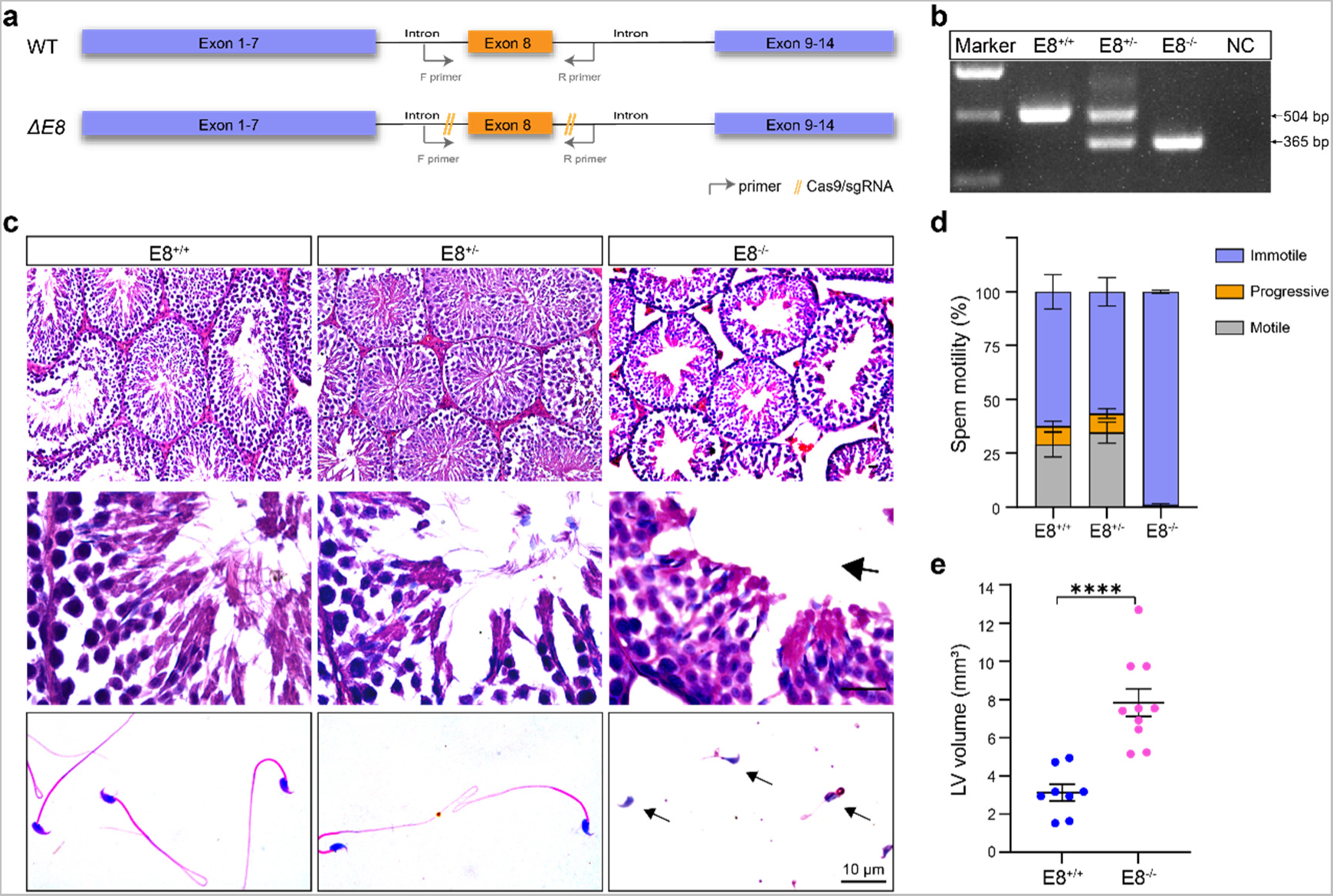
The generation and phenotypic analysis of *Katnal2 ΔE8* **mice.** **a**, Schematics of the deletion of exon 8 (*ΔE8*) of mouse *Katnal2*. **b**, PCR genotyping of *ΔE8* mice. **c**, Hematoxylin-eosin-stained testis sections (upper and middle) and Giemsa-stained cauda epididymal sperms (lower) from mice of different genotypes. **d**, Quantitative analysis of sperm motility of different genotypes. Error bars are s.e.m. **e**, Quantitative analysis of lateral ventricle (LV) volumes of WT and *ΔE8* mice. **** *p* < 0.0001, Student’s *t*-test.

**Extended Data Fig 3.**
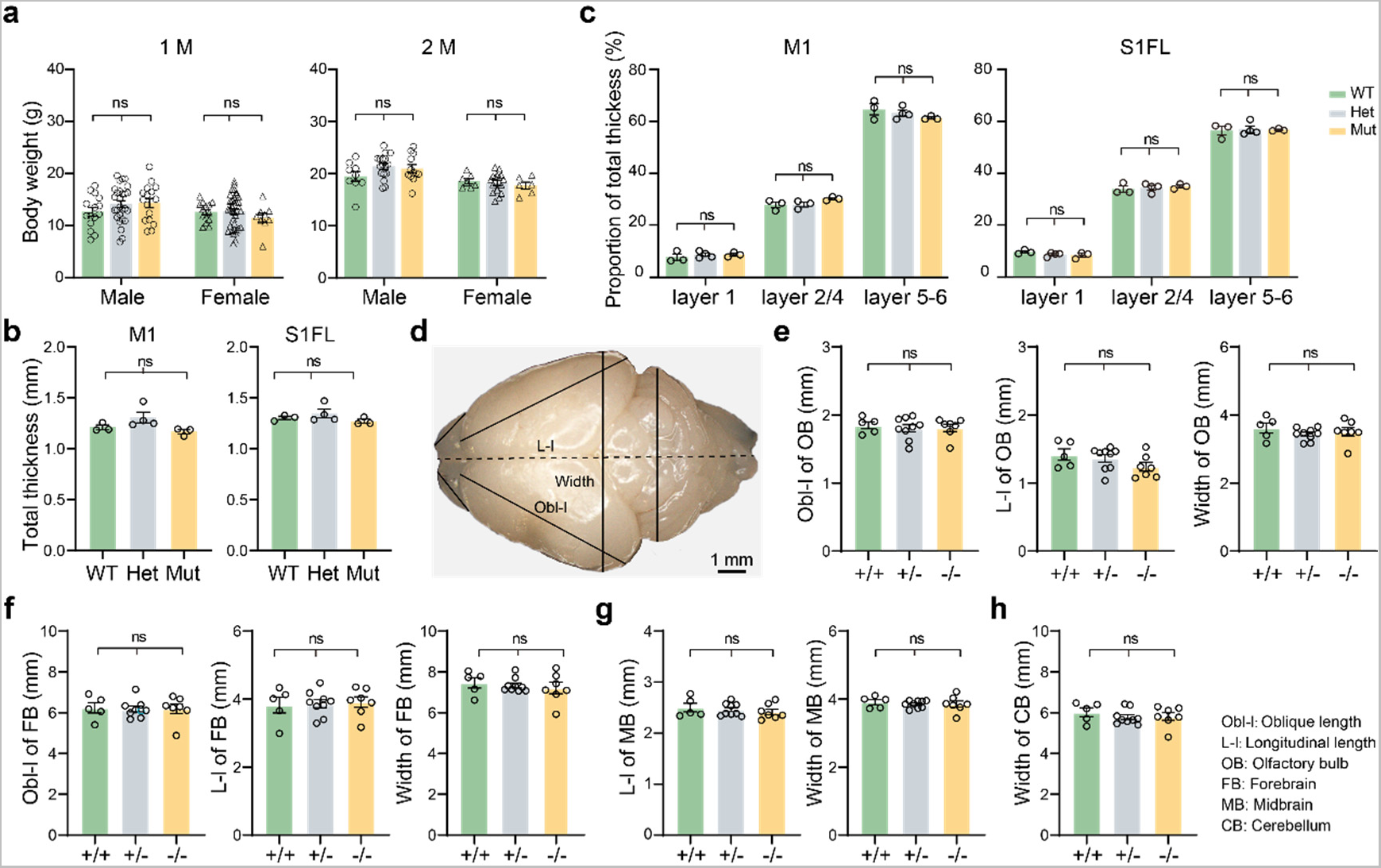
Morphological analysis of *Katnal2* mutant mice. **a**, Comparison of the body weight of 1- and 2-month-old mice of different genotypes. **b**, Comparison of the thickness of M1 and S1FL cortex in 2-month-old mice of different genotypes. M1: primary motor cortex; S1FL: primary somatosensory cortex forelimb region. **c**, Comparison of the relative thickness of different layers of M1 and S1FL cortex among WT, Het, and Mut mice. **d**, Morphological measurement of mouse brains. Black straight lines and dashed lines indicate different measurements. **e-f**, Comparison of the oblique length (Obl-l), longitudinal length (L-l) and width of the olfactory bulb (OB, **e**) and the forebrain (FB, **f**) among mice of different genotypes. **g**, Comparison of the longitudinal length and width of the midbrain (MB) among mice of different genotypes. **h**, Comparison of the width of the cerebellum (CB) among mice of different genotypes. Data are presented as mean ± s.e.m. Each data point represents the result from one animal. ns: not significant, one-way ANOVA test.

**Extended Data Fig 4.**
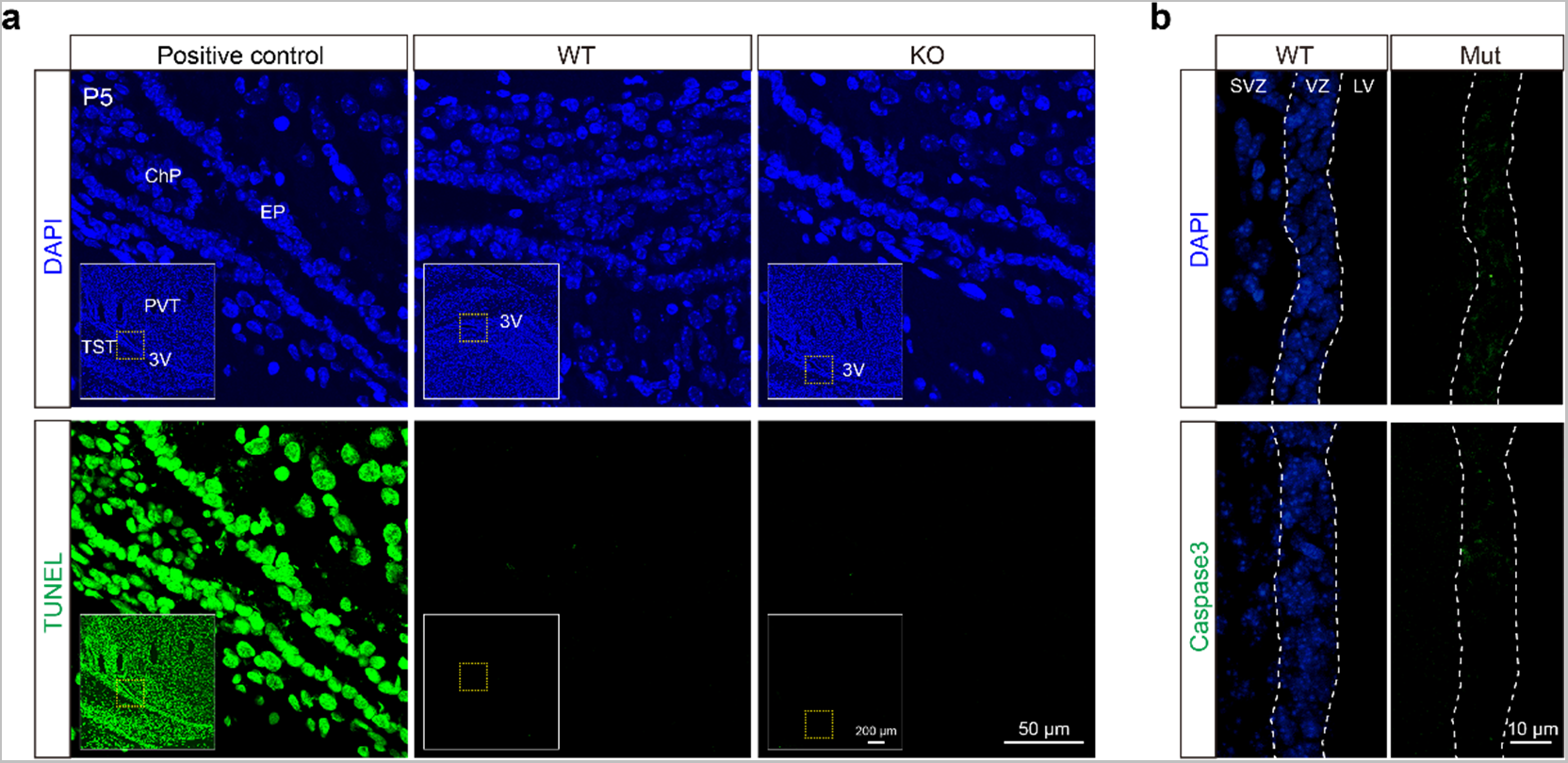
*Katnal2* gene deficiency does not cause massive cell apoptosis in the cortex of neonatal mice. **a**, *Katnal2* gene knockout (-/-) did not increase TUNEL^+^ cells (green) in the VZ/SVZ region of the cortex at P5. Positive control of TUNEL staining was conducted using brain slices from WT littermates. Ep, ependyma; PVT, paraventricular thalamic nucleus; TST, tectospinal tract. High-magnification images are from the selected areas of low-magnification images of the brain slice shown in the inserts. **b**, *Katnal2* mutation did not cause an increase in the cleaved-Caspase 3^+^ cells.

**Extended Data Fig 5.**
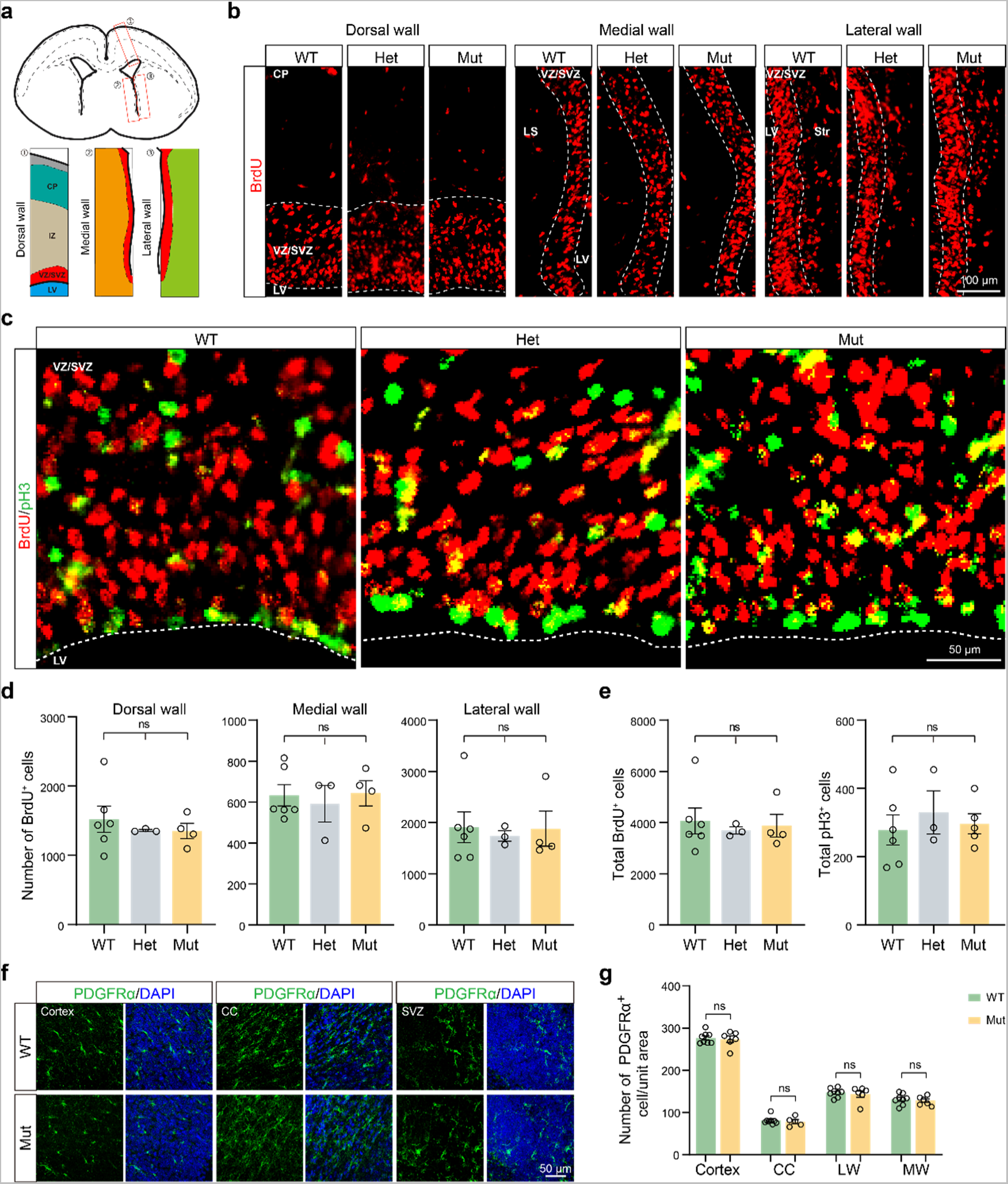
*Katnal2* deficiency does not impair neuro-gliogenesis genesis. **a**, Schematics of the dorsal, medial, and lateral walls of the LV. IZ, intermediate zone; LS, lateral septum; Str, striatum. **b,** Fluorescent microscopic graphs showing the 24-hr BrdU incorporation in the VZ/SVZ of different regions (dorsal wall, lateral wall, and medial wall) at the anterior LV in E17 mice of different genotypes. Dashed lines outline the VZ/SVZ area. **c**, Fluorescent microscopic graphs showing the co-staining of BrdU and phosphor-Histon3 (pH3) at the VZ/SVZ of the dorsal wall of the LV in WT, Het, and Mut mice at E17. Dashed lines outline the edge of the LV. **d**, Quantitative analysis of BrdU^+^ cells in the VZ/SVZ of the dorsal wall, lateral wall, and medial wall of the LV from WT, Het, and Mut mice at E17.5. **e**, Quantitative analysis of total BrdU^+^ cells and pH3^+^ cells in the VZ/SVZ of the LV at E17.5. **f-g**, *Katnal2* deficiency did not alter the density of oligodendrocyte precursors. PDGFR ^+^ oligodendrocyte precursor cells (OPCs) in different brain regions were labeled by immunofluorescent staining. CC, corpus callosum; MW, SVZ of the medial wall of the LV; LW, SVZ of the lateral wall of the LV. Quantitative data are presented as mean ± s.e.m. ns: not significant, one-way ANOVA.

**Extended Data Fig 6.**
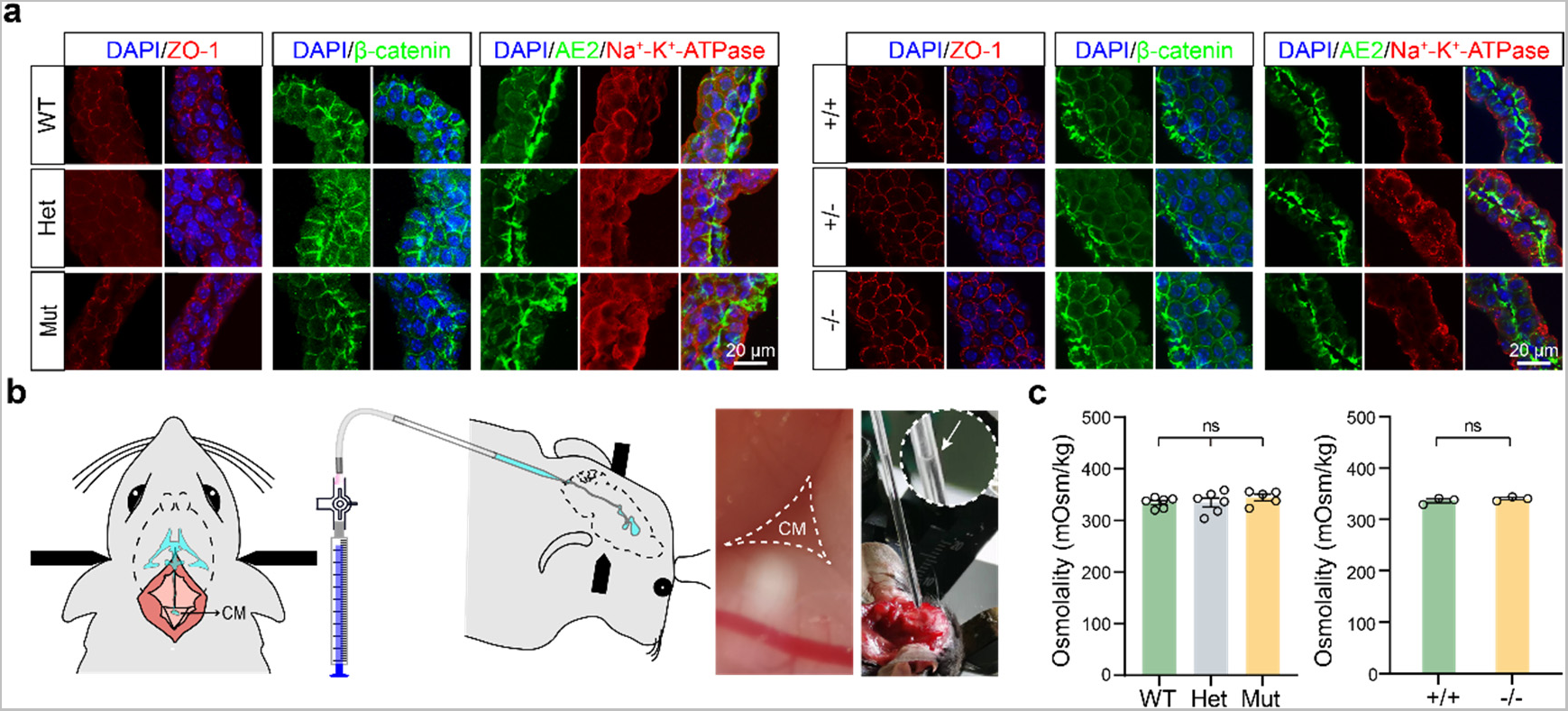
Distribution of functional proteins in the choroid plexus and osmotic pressure of the cerebral spinal fluid are not affected by *Katnal2* deficiency. **a**, Immunofluorescent analysis of the distribution of ZO-1, β-catenin, AE2, and Na^+^-K^+^-ATPase in the ChP of *Katnal2* Mut mice and *Katnal2* KO mice. **b**, A schematic diagram and photograph of the aspiration of cerebrospinal fluid (CSF) from the cisterna magna (CM) of adult mice. **c**, Osmolality of CSF from *Katnal2* Mut mice and *Katnal2* KO mice compared to their WT or Het littermates. Each data point represents the result from one animal. ns: not significant, one-way ANOVA test.

**Extended Data Fig 7.**
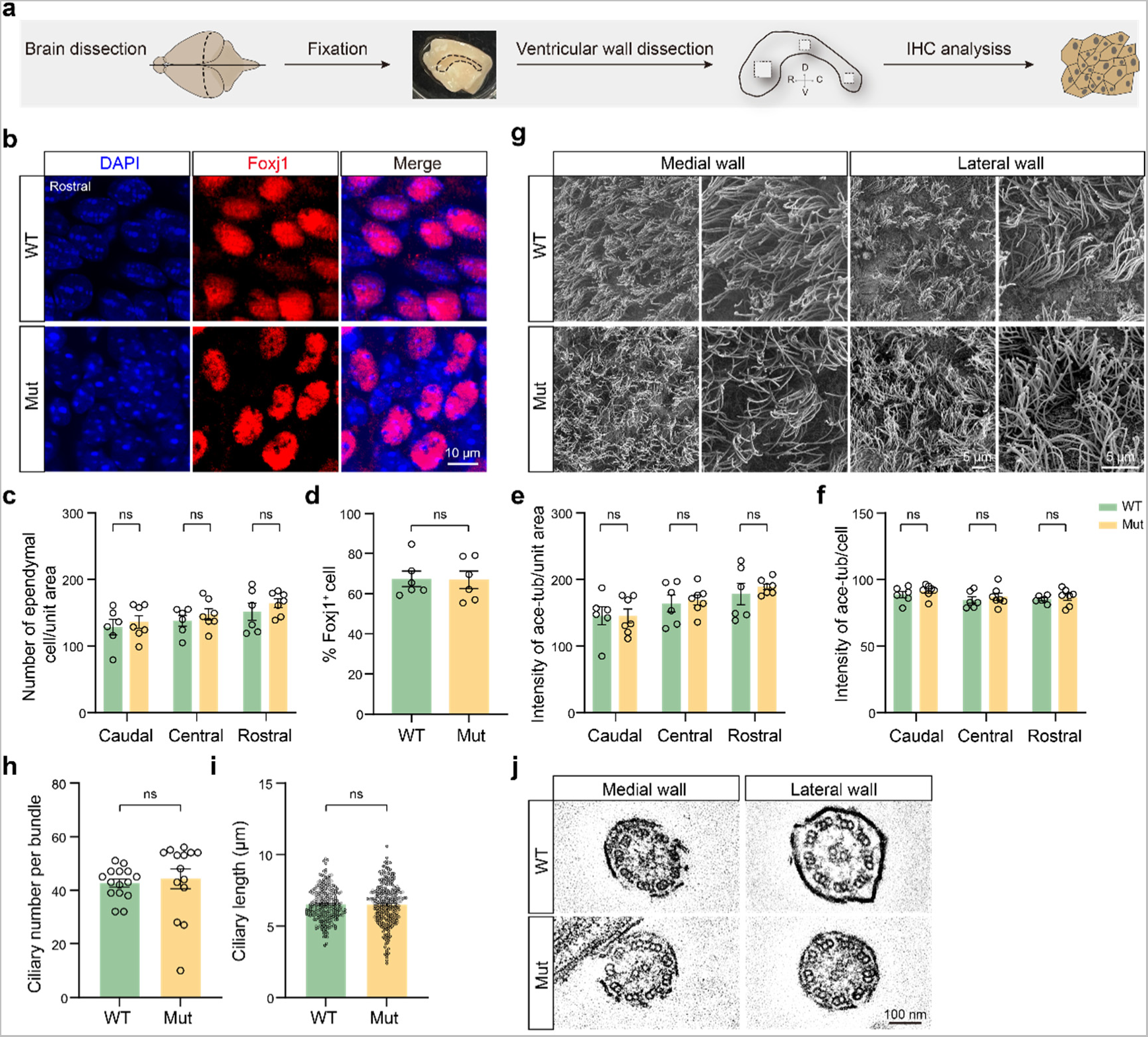
Normal ependymal cell maturation and cilium structure in *Katnal2* mutant mice. **a**, Schematic diagram of immunohistochemistry in tissue samples of the lateral wall of the LV. **b**, Immunofluorescent staining of Foxj1 (for mature EpCs) in the rostral region of the lateral wall of the LV in WT and Mut mice at 2 months of age. **c-h**, There is no significant difference between Mut mice and WT littermates in all following parameters: EpC number per unit area (**c**), the proportion of Foxj1^+^cells among EpCs at the lateral wall of the LV (**d**), the fluorescent intensity of acetylated tubulin (ace-tub) per unit area (**e**), ace-tub fluorescent intensity per EpC (**f**), cilium number per bundle (**h**), and the mean ciliary length of EpCs (**i**). Each data point represents the result from one animal in **c-f**, from one EpC in **h**, and from one cilium in **i.** WT: n = 256 cilia of 15 cilium bundles from 3 mice; Mut: n = 278 cilia of 14 cilium bundles from 3 mice in **h** and **i**. **g,** Scan electron microscopic images of ependymal cilia on the medial and lateral walls of the LV in WT and Mut mice. Cilia in each bundle were more scattered in Mut mice, while well-aligned in WT mice. **j**, Transmission electron microscopic images showing the normal microtubule organization of individual ependymal cilia on the medial and lateral walls of the LV in Mut mice.

**Extended Data Fig 8.**
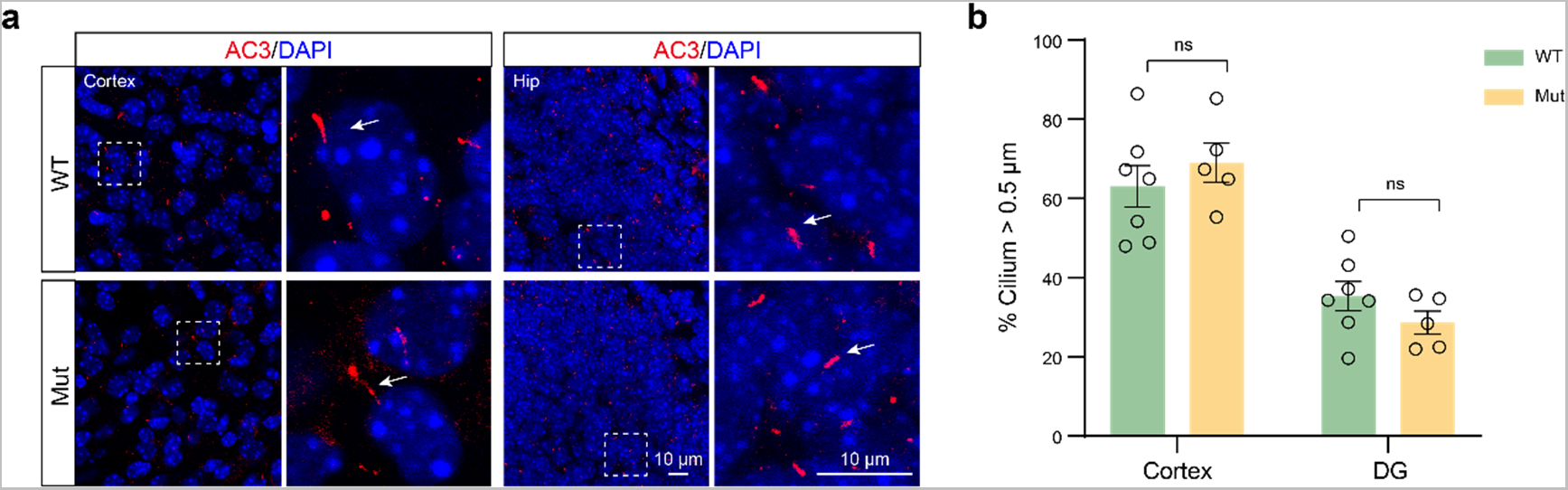
Normal morphogenesis of primary cilia in the cerebral cortex and hippocampus of *Katnal2* mutant mice. **a**, Immunofluorescent staining of primary cilia in the cortex and hippocampus of WT and Mut mice using the ciliary marker adenylate cyclase 3 (AC3). **b**, Quantitative analysis of the proportion of cells with the cilium length > 0.5 mm in the cortex and hippocampus. ns: not significant, one-way ANOVA test.

**Extended Data Fig 9.**
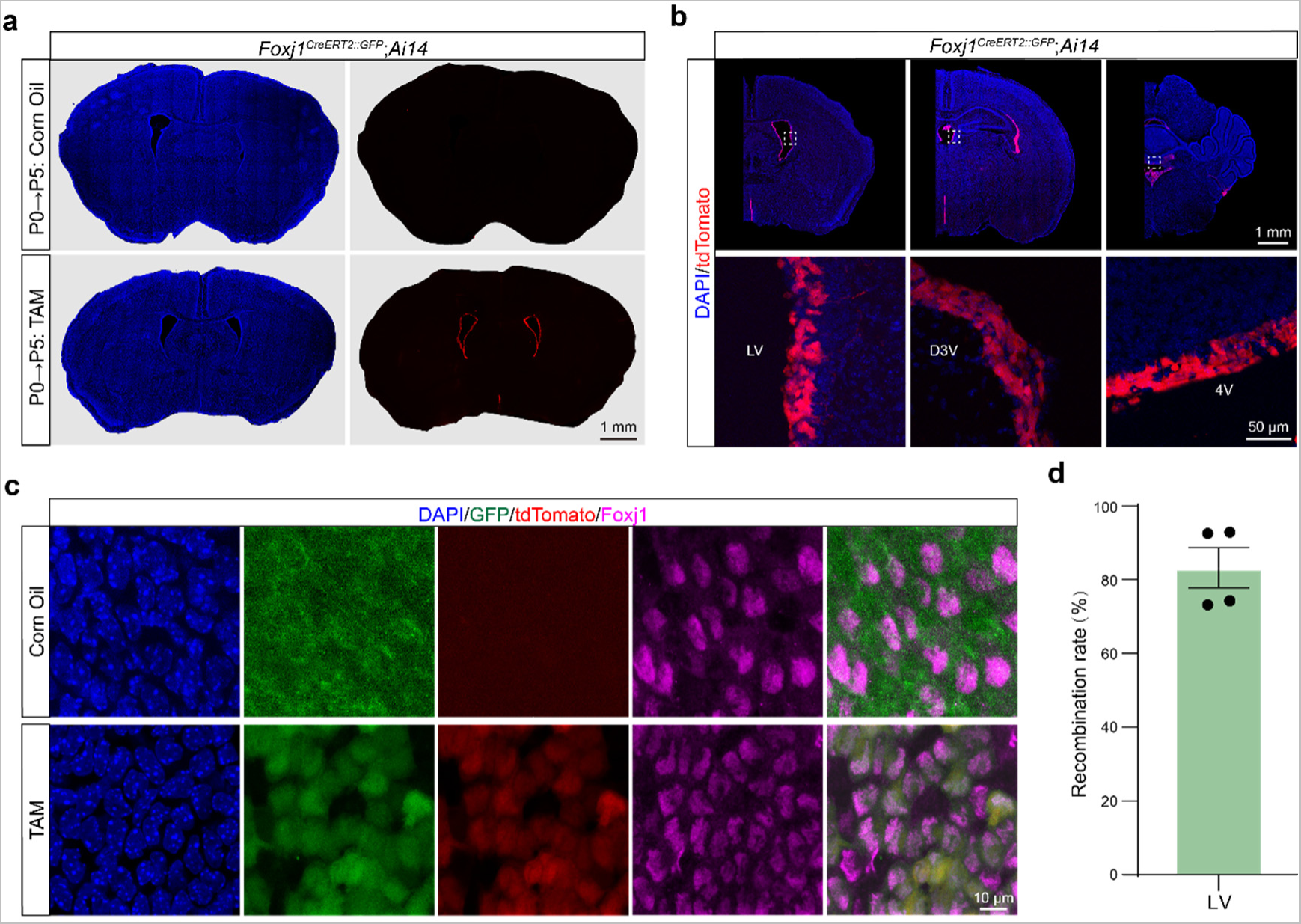
Validation of ependyma-specific recombination of the conditional allele by *Foxj1^CreER^*. **a**, Validation of the induction of *Foxj1^CreER^-*mediated recombination in newborn mice by maternal application of tamoxifen (TAM) daily from P0 to P5. The Ai14 fluorescent reporter line was used to indicate the Cre-mediated recombination (red). Corn oil was administrated as the negative control. Note: there is no leakage of Cre recombinase activity in the absence of TAM. **b**, Validation of *Foxj1^CreER^-*mediated recombination in EpCs of all four brain ventricles upon maternal application of TAM. **c**, Immunofluorescent staining showing *Foxj1^CreER^*-mediated recombination specifically in Foxj1^+^ EpCs at the LV wall after maternal application of TAM. **d**, The induction rate of *Foxj1^CreER^*-mediated recombination in EpCs the LV wall of newborn mice by maternal application of TAM. Each data point represents the result from one animal.

**Extended Data Table 1. Gene co-expression analysis of *KATNAL2*.**

**Extended Data Table 2. Disease-associated mutations in the coding region of *KATNAL2*.**

**Extended Data Table 3. scRNA-seq analysis of P5 mouse cortex.**

**Extended Data Table 4. Antibodies used for immunofluorescent staining.**

**Extended Data Movie 1. Ependymal flow of adult WT and Mut mice indicated by the flow of fluorescent beads in an acute whole-mount preparation of lateral ventricles.**

**Extended Data Movie 2. Phase-contrast time-lapse microscopy showing the beating of ependymal cilia in brain slices from adult WT and Mut mice.**

**Extended Data Movie 3. Beating of ependymal cilia of adult WT and Mut mice indicated by the motion of fluorescent beads attached to the cilium tips.**

## Supplementary Information

**A summary of data analysis and statistics in this study.**

